# βII and βIII spectrin paralogues define robustness and specialization of the neuronal membrane periodic skeleton

**DOI:** 10.64898/2026.03.23.713115

**Authors:** Marie-Lise Jobin, Léa Sarzynski, Magali Mondin, Tatiana Charbonnier, Sophie Daburon, Nicolas Chevrier, Pauline Belzanne, Isabelle Jansen, Evelyn Garlick, Matthieu Sainlos, Daniel Choquet, Anna Brachet

## Abstract

Neuron connectivity and signal processing rely on a complex, dynamic morphology regulated by the cytoskeleton. The membrane-associated periodic skeleton (MPS), a 190-nm periodic actin-spectrin lattice, is a conserved feature of neurons. Here, we show that the dendritic MPS is built from a dual β-spectrin system in which βII- and βIII-spectrins are co-expressed and interleaved at the nanoscale in shafts and spine necks. 3D MINFLUX nanoscopy suggests that these paralogues form both homotypic (βII- or βIII-only) and heterotypic (βII/βIII) tetramers with an approximately 100-nm radial periodicity. Either paralogue alone is sufficient to maintain lattice architecture, whereas their combined loss disrupts MPS integrity. Using targeted mutagenesis, we show that actin binding is required to stabilize both paralogues within the MPS, whereas βIII-spectrin additionally depends on phosphoinositide interactions. Our findings reveal that, unlike the axonal MPS, the dendritic MPS is a composite scaffold in which structural redundancy coexists with paralogue-specific regulatory mechanisms, with potential consequences for synaptic function.

## Introduction

Neurons exhibit a highly polarized morphology consisting of a cell body, a long and thin axon, branched dendrites and thousands of synaptic connections. This architecture, essential for neuronal computation and connectivity, is built and dynamically remodeled by the neuronal cytoskeleton^1^. Among the cytoskeletal assemblies, the membrane-associated periodic skeleton (MPS) is a unique feature of neural cells^2–4^ and a peculiar cellular organization discovered a decade ago by super-resolution microscopy^5^. The MPS consists of circumferential actin rings interconnected by spectrin tetramers at ∼190 nm intervals beneath the plasma membrane^6^. First identified in axons^5^, the MPS was later visualized in a subset of dendrites and dendritic spine necks^7–10^. In axons, the MPS governs structural integrity^11,12^ and physiology by controlling axonal caliber^13,14^, organizing ion channels^15–17^ and signaling complexes^18,19^, and regulating membrane trafficking and plasticity at the axon initial segment (AIS)^20^. Although the role of the axonal MPS is beginning to emerge, its precise nanoscale organization, molecular composition, and function in dendritic compartments remain poorly understood. Addressing this gap is essential as dendrites, and especially dendritic spines, must simultaneously maintain long-term structural stability and support rapid, activity-dependent morphological remodeling, two processes that rely heavily on the regulation of their actin cytoskeleton^21^. Whether the periodic actin-spectrin scaffold contributes to robustness, plasticity, or both in dendritic compartments remains unclear.

Spectrins are ancient cytoskeletal elements that emerged early in metazoan evolution as single α- and β-spectrin genes, providing basic membrane support^22–24^. Gene duplication and divergence during vertebrate evolution expanded the spectrin family, giving rise to several paralogues with specialized localizations and roles^22–24^. Among β-spectrins, βII-spectrin is broadly expressed and displays compartment-specific segregation within individual cells, where it organizes specialized domains, such as the transverse tubules in cardiomyocytes^25^ or the lateral membranes in epithelial cells^26^. βIV-spectrin localizes to and organizes excitable membrane regions, including the AIS in neurons^17,27,28^ and the intercalated disc in cardiomyocytes^29^. βIII-spectrin, the most recent paralogue and restricted to mammals^24^, is expressed in the pancreas, kidney, and several brain regions^30,31^, including the cerebellum and hippocampus. In neurons, αII-spectrin pairs with compartment-specific β-spectrin paralogues^32,33^: βII-spectrin in the soma, axon, dendrites, and spines; β4-spectrin at the AIS and nodes of Ranvier; and βIII-spectrin in the somatodendritic compartment^34^ (Fig. 1a).

**Figure 1.**
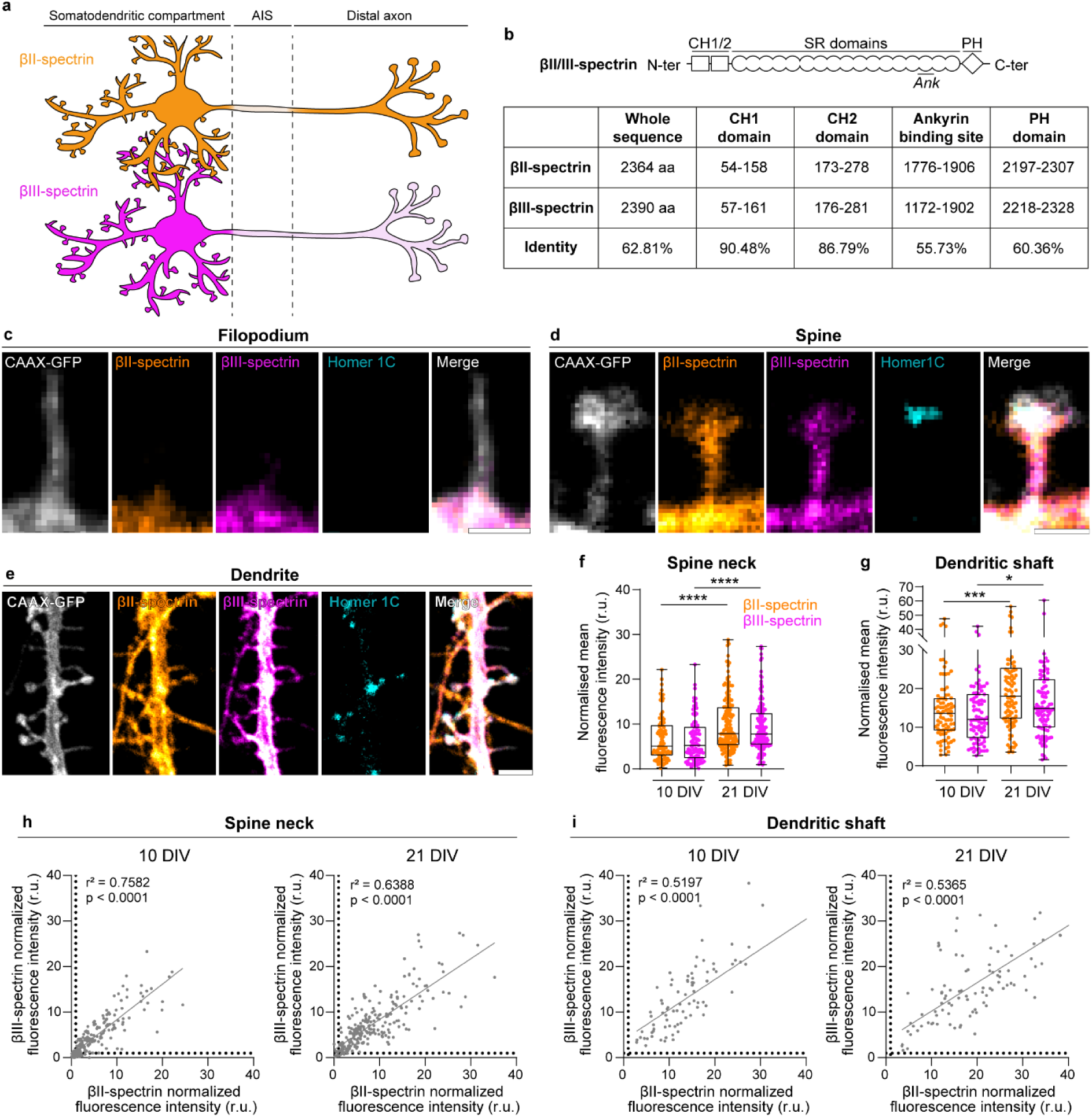
βII- and βIII-spectrins are co-expressed in dendrites and dendritic spine necks. **a,** Schematic of βII and βIII spectrins expression in the somatodendritic compartment, axon initial segment (AIS) and distal unmyelinated axon. **b,** Comparison of βII and βIII spectrins sequences and binding domains. **c, d, e,** Representative confocal images for βII-spectrin (orange), βIII-spectrin (magenta), CAAX-GFP (grey) and Homer1C (cyan) in a filopodium (a), a mature spine (b) or a dendrite (c) at 21 DIV. Scale bars, 1 µm for filopodium and spine, 2 µm for dendrite. Note that the display contrast was purposely kept constant. **f, g,** Fluorescence intensity of βII-spectrin and βIII-spectrin in spine necks and dendrites at 10 DIV and 21 DIV normalized to the average intensity of filopodia at the same time-point. Each box shows the median ± percentile. n between 98 and 160 spines or 84 and 97 dendrites from 20 to 30 neurons examined over 3 independent experiments for each time point. Mann-Whitney test, *: p < 0.05, ***: p < 0.001, ****: p < 0.0001. **h, i,** Correlation scatter plots between βII-spectrin and βIII-spectrin normalized fluorescence intensities in spine neck or dendrite at 10 DIV and 21 DIV. n between 98 and 160 spines or 84 and 97 dendrites from 20 to 30 neurons examined over 3 independent experiments for each time point. r^2^, Pearson’s r coefficient; P, P value.

βII- and βIII- spectrins share a similar domain organization^23^, including 17 spectrin repeats (SRs), N-terminal tandem calponin homology (CH) domains mediating interaction with F-actin^35^, an ankyrin-binding site within SR14–15^36^, and a C-terminal pleckstrin homology (PH) domain that interacts with membrane phosphoinositides^37^ (Fig. 1b and S1). While the CH domains are highly conserved, the ankyrin-binding and PH regions show lower sequence homology (Fig. 1b and S1), raising the possibility that the two paralogues may be differentially regulated.

Dendrites and dendritic spines represent a rare example among vertebrate subcellular compartments, and a unique case within neuronal domains, in which two β-spectrin paralogues coexist (Fig.1a), raising the question of whether they play specialized roles or act redundantly to build and regulate the dendritic MPS.

Here, we address the organization and function of βII- and βIII-spectrins in dendritic shafts and spines by combining advanced fluorescence nanoscopy, CRISPR-mediated knockout, targeted mutagenesis, and live imaging in hippocampal neurons. We show that βII- and βIII-spectrins are co-expressed in all dendritic shafts and spine necks and are interleaved within the MPS lattice at the nanoscale. 3D-MINFLUX imaging suggests both homotypic (βII- or βIII-only) and heterotypic (βII/βIII) tetramers arranged in a radially periodic organization. Selective depletion of either paralogue leaves MPS architecture largely intact, whereas dual removal abolishes it, demonstrating that the two isoforms act as redundant structural units of the dendritic MPS. Finally, targeted mutagenesis combined with super-resolution and live imaging demonstrate that actin binding is essential for the stabilization and MPS incorporation of both paralogues, while βIII-spectrin uniquely relies on phosphoinositide interaction. Together, these findings demonstrate that β-spectrin paralogues act as redundant yet differentially regulated scaffolds conferring both robustness and adaptive flexibility to the MPS in dendritic shafts and spines.

## Results

### βII and βIII spectrins are co-expressed in dendritic shafts and spine necks

To determine whether βII- and βIII-spectrins occupy strictly overlapping or distinct subcellular domains, we first systematically examined their spatial distribution in dendrites.

Primary dissociated hippocampal neuronal cultures were transfected with a membrane-targeted GFP (CAAX-GFP) to visualize dendritic morphology and spines. Cells were fixed at two developmental stages, 10 and 21 days in vitro (DIV), and immunolabeled for endogenous βII- and βIII-spectrins together with Homer1C, a marker of the mature postsynaptic compartment (Fig. 1). Based on these markers, dendrites were distinguished from axons, and spines were classified as either immature (filopodia), which lacked Homer1C labeling (Fig. 1c), or mature, which exhibited a distinct neck and a Homer1C-positive head (Fig. 1d). Fluorescence bleed-through between channels was negligible, with 0% from the βIII to the βII-spectrin channel and 0.38% from the βII to the βIII-spectrin channel (Fig. S2). Consistent with previous reports, βIII-spectrin was largely absent from axons^30,38,39^, and analysis of this paralogue was therefore restricted to dendritic shafts and spines. Filopodia contained very low levels of βII- and βIII-spectrins at 10 and 21 DIV, whereas most dendritic shafts and mature spines contained both paralogues (Fig. 1c–e). In spines, βII- and βIII-spectrins localized primarily to the neck and the base of the head (Fig. 1c). Quantitative analysis showed that β-spectrin signal intensities were approximately 5-fold higher in spine necks and 12–15-fold higher in dendritic shafts compared with filopodia at 10 DIV, increasing to 8-fold and 15–20-fold enrichment, respectively, at 21 DIV (Fig. 1f, g). βII- and βIII-spectrin signals were strongly correlated in both dendritic shafts and spines, indicating that the two paralogues are consistently co-expressed within the same compartments across developmental stages (Fig. 1h, i).

Together, these data indicate that βII- and βIII-spectrins are largely absent from filopodia but are co-expressed and spatially correlated in most dendritic shafts and mature spines.

### βII- and βIII-spectrins are interleaved within the MPS in dendritic shafts and spines

As no distinct subpopulation of dendritic shafts or spines preferentially expressed βII- or βIII-spectrin, we next examined their nanoscale co-organization within these compartments with different super-resolution imaging modalities.

Using STochastic Optical Reconstruction Microscopy (*d*STORM)^40^, we quantified the periodic organization of βII-spectrin in axons and of both βII- and βIII-spectrins in dendritic shafts and spines of mature neurons (21 DIV). Neurons expressing cytosolic mCherry served as a non-periodic control. Dendritic spines were identified based on morphology and Homer1C or PSD95 staining, whereas neurites lacking spines were classified as axons (Fig. 2). Spatial periodicity in line-scan intensity profiles was quantified as the amplitude at the first peak of the autocorrelation function, which reports how strongly the signal repeats when shifted by one spatial period. Validation of the imaging and analysis pipeline using βII-spectrin in axons confirmed robust periodicity (i.e. high autocorrelation amplitudes) and a spacing of ∼188 nm, consistent with previous reports^3,5^ (Fig. 2a, d). In dendritic shafts and spine necks, autocorrelation values were lower than in axons but remained significantly above non-periodic controls, with spacings comparable to those measured in axons (Fig. 2b–d). Applying a threshold defined by the mean autocorrelation amplitude of cytosolic mCherry, 98% of axons, 80% of dendritic shafts, and 64% of spines exhibited periodic βII-spectrin organization. βIII-spectrin amplitudes displayed a comparable nanoscale organization, with a spacing of ∼180 nm in dendritic shafts and spine necks (Fig. 2g). Its autocorrelation amplitudes (Fig. 2e, f) and the fraction of periodic structures (68% of dendritic shafts and 62% of spines) were similar to βII-spectrin and significantly above non-periodic controls, indicating that indeed βIII-spectrin also forms a periodic lattice in these compartments.

**Figure 2.**
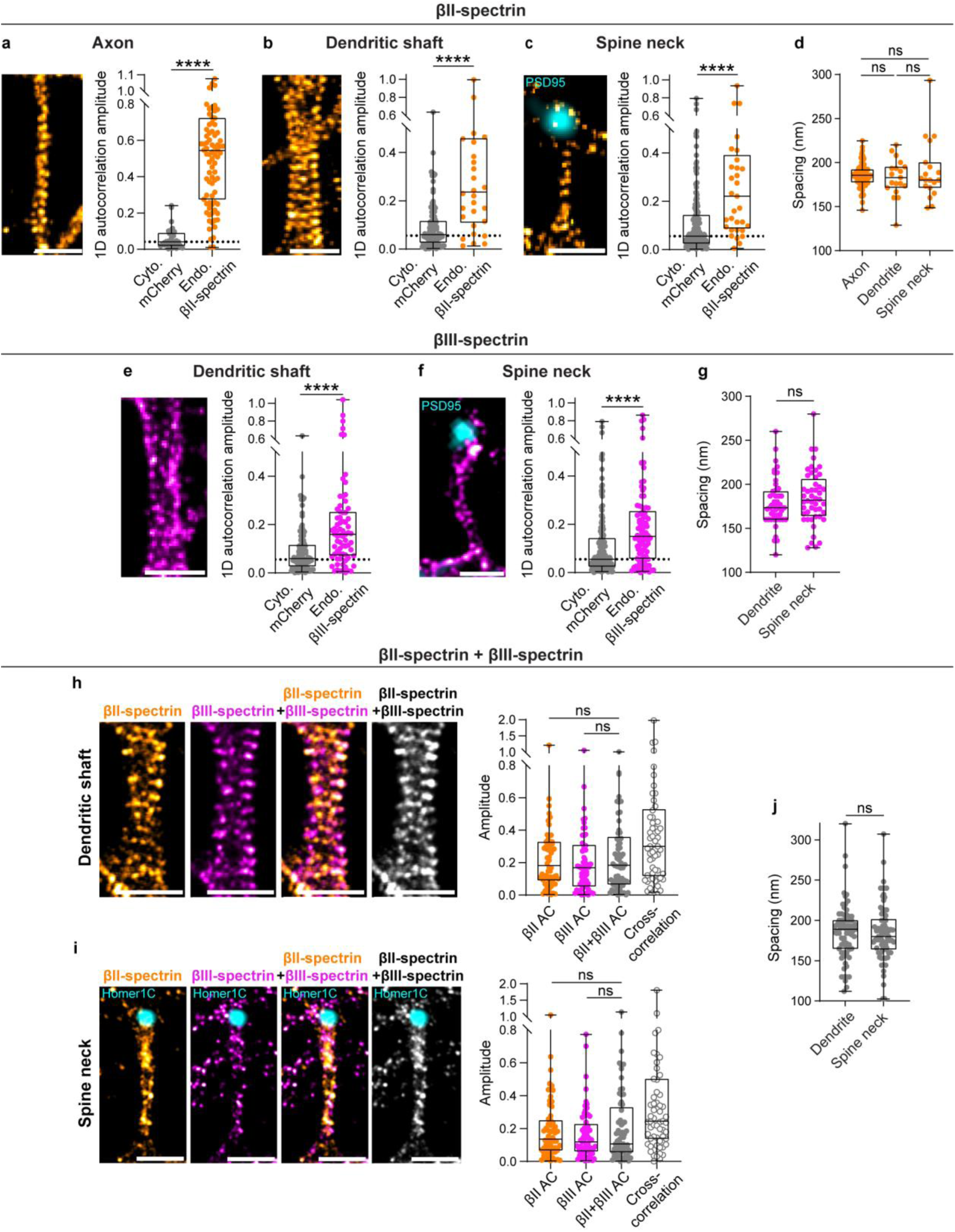
βII- and βIII-spectrins are interleaved in the MPS lattice in dendrites and dendritic spine necks. **a, b, c,** Representative *d*STORM images of βII-spectrin and its 1D autocorrelation amplitude in axons (a), dendrites (b), and spine necks with PSD95 labelling observed in epifluorescence (c). The non-periodic control was obtained by measuring the 1D autocorrelation amplitude of a cytosolic mCherry labelling in axons, dendrites or spine necks. n between 45 and 80 regions of interest examined over at least 3 independent experiments. **d,** Spacing between βII-spectrin epitopes. **e, f,** Representative *d*STORM images of βIII-spectrin and its 1D autocorrelation amplitude in dendrites (e) and spine necks (f). n between 45 and 80 regions of interest examined over at least 3 independent experiments. **g,** Spacing between βIII-spectrin epitopes. **h, i,** Representative two-color STORM images of βII-spectrin and βIII-spectrin and autocorrelation (AC) or cross-correlation amplitudes in dendrites (h) and spine necks with Homer1C labelling (i). n = 76 regions of interest examined over 3 independent experiments. **j,** Spacing between β-spectrin epitopes after merging the two channels. Each box shows the median ± percentile. Scale bars, 1 µm. Mann-Whitney test, ns: non-significant, ****: p < 0.0001.

To assess the co-organization of βII- and βIII-spectrins at the nanoscale within the MPS in dendritic shafts and spines, we performed two-color *d*STORM super-resolution. Under dual labeling, the periodic organization of both βII- and βIII-spectrins was comparable to that observed in single-color acquisitions, in both spacing and autocorrelation amplitude (Fig. 2h–j), indicating that two-color immunolabeling did not interfere with epitope detection. Merging the two fluorescence channels did not increase autocorrelation amplitude relative to either channel alone (Fig. 2h, i), suggesting that βII- and βIII-spectrins are sufficiently abundant to populate all periodic units when analyzed individually. We then quantified the relative organization of βII- and βIII-spectrin signals using cross-correlation, which measures their similarity as a function of spatial lag. The cross-correlation analysis, quantified as the amplitude at the first period, revealed intermediate values of 0.2–0.3 in dendritic shafts and spine necks (Fig. 2h, i), indicating again partial co-organization together with a shared periodic alignment of the two paralogues.

To determine the relative positioning of βII- and βIII-spectrins with nanometer precision in three dimensions, we used three-dimension MINimal emission FLUXes (3D-MINFLUX)^41^, a light microscopy technique that allows the localization of molecules with few nanometers precision^42^. In the XY plane, 3D-MINFLUX imaging revealed a punctate, dot-like pattern with more pronounced periodicity in dendrites than in spine necks (Fig. 3c, e). In dendritic shafts, independently of the β-spectrin paralogue, many localization clusters appeared in pairs, which could correspond to the two C-terminal β-spectrin epitopes present in individual spectrin tetramers (Fig. 3a and c, arrows). Using XZ projections oriented perpendicular to the dendritic axis (Fig. 3a), we resolved the radial distribution of β-spectrins (Fig. 3d, f). From this perspective, most localization clusters were also organized in pairs, and a substantial fraction contained both βII- and βIII-spectrin signals, either overlapping or immediately adjacent, suggesting that some spectrin tetramers incorporate different β-spectrin paralogues (Fig. 3d, f). Inter-cluster distance measurements, independent of paralogues identity, revealed a radial periodicity with an average spacing of approximately 100 nm in both dendrites and spine necks (Fig. 3b). Considering that individual spectrin tetramers are rod-like assemblies, with an approximate section diameter of ∼2–3 nm^43^, this organization suggests that in dendritic shafts and spines, spectrin tetramers do not form a continuous submembranous layer but instead assemble as discrete, periodically of ∼100nm-spaced elements.

**Figure 3.**
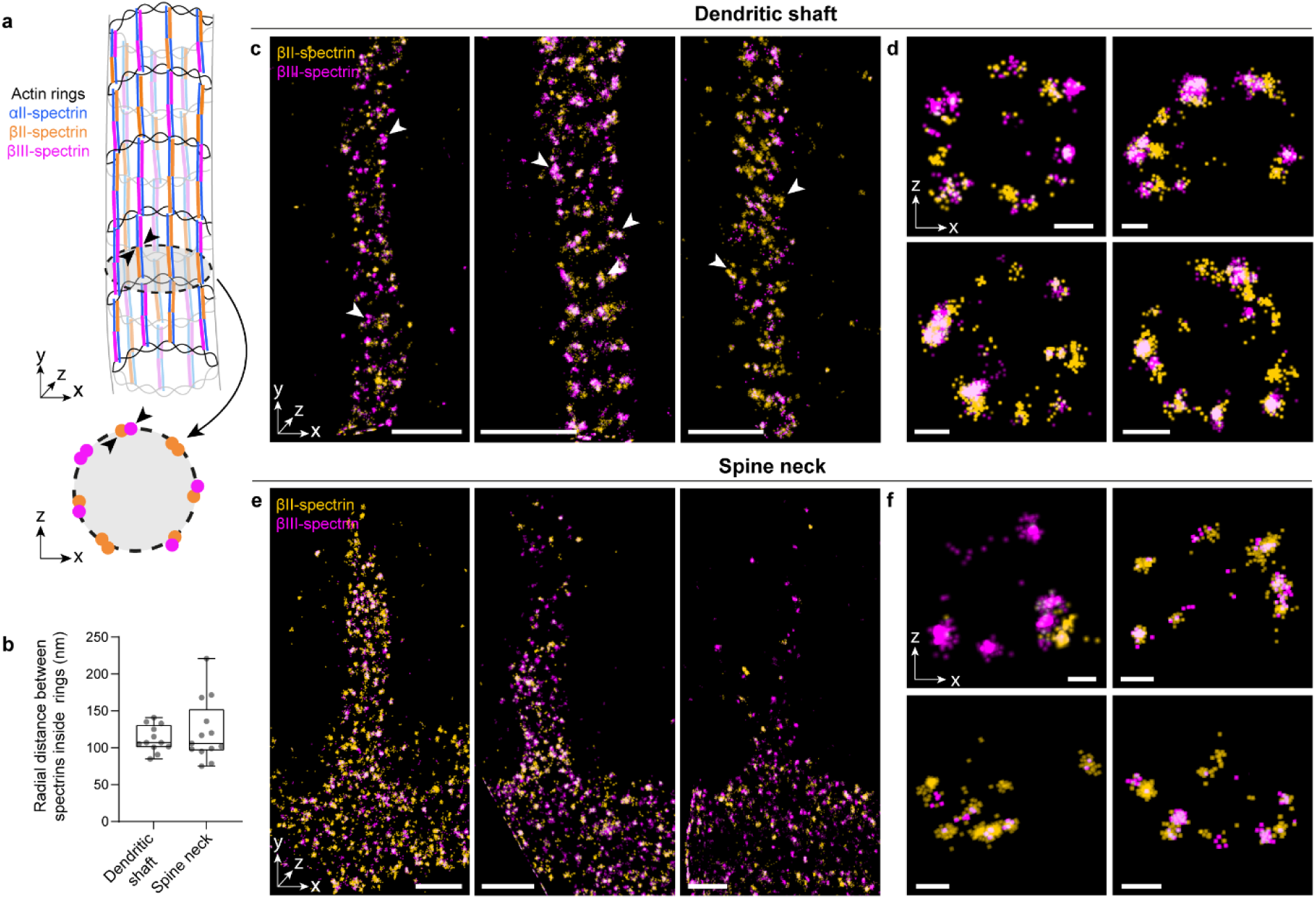
βII- and βIII-spectrins exhibit a radial periodic organization and are observed in heterotypic (βII/βIII) and homotypic (βx/βx) tetramers. **a,** Schematic of βII-, βIII- and ɑII-spectrin in the dendrite. Arrowheads correspond to the same area. **b**, Quantification of the distances between clusters of localizations on XZ projections. **c-f**, Two-color 3D MINFLUX of βII- and βIII-spectrins in dendrite (c and d) and dendritic spine neck (e and f). Scale bars, 500 nm in c and e, 100 nm in d and f. Arrowheads highlight clusters of localizations that appear in pairs, independently of the β-spectrin isoform, mostly visible in dendrites. (d) and (f) XZ projections of regions in (c) and (e) respectively, show discrete clusters regularly spaced independently of the β-spectrin paralogue, suggesting that βII- and βIII-spectrins are radially periodic.

Altogether, these results demonstrate that βII- and βIII-spectrins form a periodic lattice structure in dendritic shafts and spine necks and are intermixed within the MPS. Furthermore, 3D-MINFLUX data reveal a radial periodicity and support the existence of both homotypic (βII-or βIII-only) and heterotypic (βII/βIII) spectrin tetramers within the neuronal cytoskeleton.

### Structural robustness of the MPS lattice is ensured by functional redundancy of βII- and βIII-spectrins paralogues

To investigate the contribution of βII- and βIII-spectrins to MPS organization, we selectively depleted each paralogue and assessed its impact on the other. A CRISPR–Cas9 knockout (KO) approach was used, employing plasmids encoding SpCas9 together with either a control, non-targeting guide RNA (gRNA), a gRNA targeting βII-spectrin or βIII-spectrin. Co-expression of both gRNAs enabled simultaneous depletion of βII- and βIII-spectrins.

We first validated the efficiency of the CRISPR-mediated KO. Ten days after transfection, βII-spectrin expression was reduced by approximately 70% in neurons transfected with the βII-specific gRNA (Fig. S3a, b), and βIII-spectrin expression decreased by about 75% following transfection with the βIII-specific gRNA (Fig. S3c, d), confirming the high efficiency of the KO system. Co-expression of both gRNAs resulted in near-complete elimination of both spectrins (Fig. S3e, f). To assess the specificity of the approach and rule out compensatory effects, we measured βII- or βIII-spectrin labeling intensity after CRISPR–Cas9–mediated suppression of the other isoform. No cross-removal or compensatory upregulation was detected (Fig. S4). To minimize potential effects of β-spectrin depletion on neurite outgrowth or neuronal morphology, neurons were transfected between 8 and 10 DIV (Fig. S5). A low-efficiency calcium phosphate transfection protocol (<0.1% transfected neurons) was used to ensure that presynaptic input onto transfected cells remained largely unaltered, allowing the analysis of β-spectrin function in a largely wild-type neuronal network. Neurons were then transfected as described above, fixed at 21 DIV, immunolabeled for βII- or βIII-spectrins, and imaged using *d*STORM.

In axons, where βIII-spectrin is normally absent, deletion of βIII-spectrin had no detectable effect on βII-spectrin density or periodic organization (Fig. 4a). Strikingly, this was also true in most cellular regions, where both β-spectrins are normally present: no difference was measured in the labelling density, the autocorrelation amplitude and spacing of βII-spectrin in spine necks when the expression of βIII-spectrin was suppressed (Fig. 4c). Conversely, deletion of βII-spectrin did not alter βIII-spectrin levels or nanoscale organization in either spine necks or dendritic shafts (Fig. 4d, e). The only exception was a moderate reduction in βII-spectrin content in dendritic shafts lacking βIII-spectrin, although its periodic organization remained intact (Fig. 4b).

**Figure 4.**
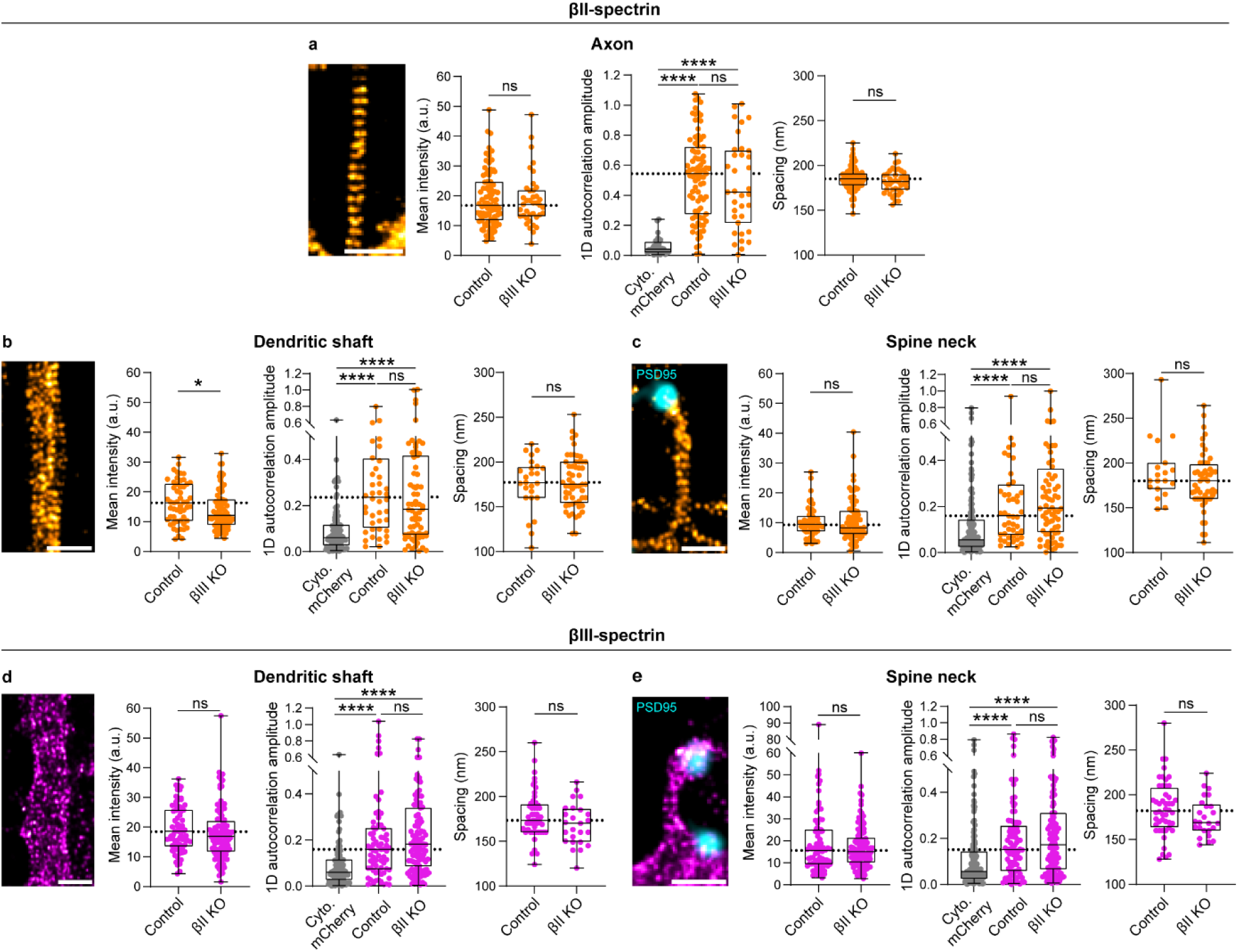
βII- and βIII-spectrins are not individually required for MPS organization. **a, b, c,** Representative *d*STORM images of βII-spectrin in a βIII-spectrin KO axon (a), dendrite (b) and spine (c), followed by βII-spectrin localization intensities, 1D autocorrelation amplitudes and spacing measured in the associated neurites. The non-periodic controls are the same as in figure 2. PSD95 was imaged using epifluorescence microscopy. n between 30 and 88 regions of interest per neurite examined over 5 independent experiments. **d, e,** Representative *d*STORM images of βIII-spectrin in a βII-spectrin KO dendrite (d) and spine (e), followed by βIII-spectrin localizations intensities, 1D autocorrelation amplitudes and spacing measured in the associated neurites. n between 30 and 88 regions of interest per neurite examined over at least 3 independent experiments. Each box represents the median with percentile. Scale bars, 1 µm. Mann-Whitney tests, ns: non-significant, *: p < 0.05, ****: p < 0.0001.

Together, these data indicate that neither βII- nor βIII-spectrin is individually required for MPS organization. This absence of effect is not explained by compensatory upregulation, as the total abundance of each paralogue remained unchanged under knockout conditions.

To further assess the relative contributions of βII- and βIII-spectrin tetramers to the structural robustness of the MPS, we examined αII-spectrin, the obligatory binding partner of β-spectrins in neuronal spectrin tetramers. We reasoned that, in the absence of one β-spectrin, the αII-spectrin signal should reflect the abundance of the remaining β-spectrin paralogue.

In dendritic shafts and spine necks, loss of either βII- or βIII-spectrin produced a comparable reduction in αII-spectrin signal (Fig. 5b, c). Simultaneous deletion of both β-spectrins resulted in an almost complete loss of αII-spectrin labeling (Fig. 5b–c). These results indicate that βII-and βIII-spectrins contribute equally to the overall spectrin tetramer content in dendritic shafts and spines, and further confirm the absence of compensatory upregulation between paralogues. Spatial organization was then analyzed on the conditions were αII-spectrin was still detected (see captions). Notably, when only one β-spectrin paralogue was present, the remaining αII-spectrin largely preserved its periodic nanoscale organization (Fig. 5b, c), suggesting that the MPS lattice remains periodically organized even when approximately half of its spectrin tetramers are lost.

**Figure 5.**
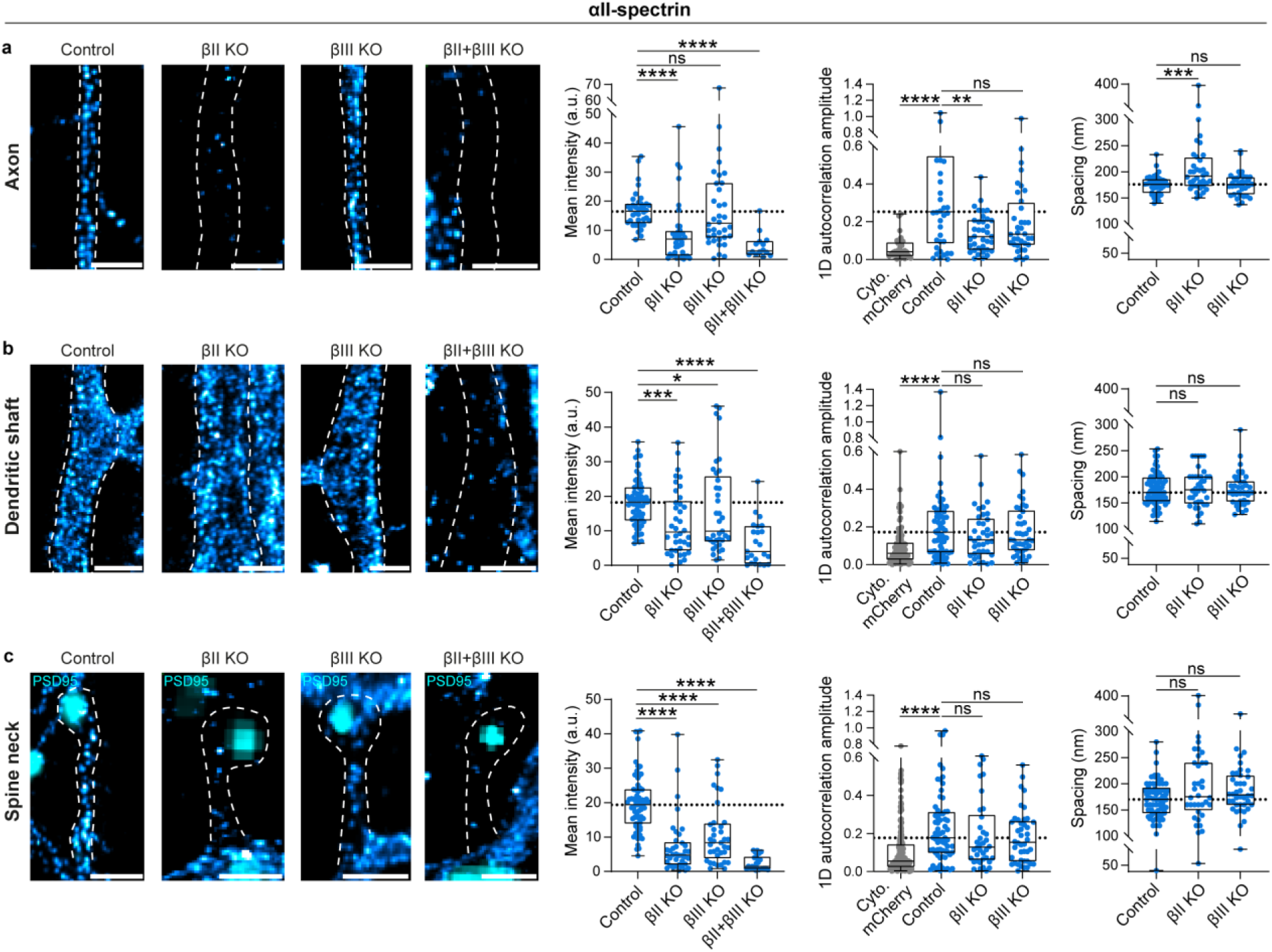
βII- and βIII-spectrins contribute equally to spectrin tetramer content and structuration in dendrites and dendritic spines. **a, b, c,** Representative *d*STORM images of ɑII-spectrin in control or KOs axons (a), dendrites (b) and spines (c), followed by ɑII-spectrin mean intensities, 1D autocorrelation amplitudes and spacing measured in the corresponding compartments. Only conditions where ɑII-spectrin mean intensity median value was above one-quarter of the controls ɑ2-spectrin values were analysed for autocorrelation and spacing. The non-periodic controls are the same as figure 2. PSD95 was imaged using epifluorescence microscopy. n between 14 and 53 regions of interest per neurite examined over 3 independent experiments. Each box represents the median with percentile. Scale bars, 1 µm. Mann-Whitney tests, ns: non-significant, *: p < 0.05, **: p < 0.01, ***: p < 0.001, ****: p < 0.0001.

In axons, as expected, deletion of βIII-spectrin did not alter αII-spectrin labeling density or spatial organization, consistent with the absence of βIII-spectrin in this compartment (Fig. 5a). In contrast, removal of βII-spectrin, either alone or in combination with βIII-spectrin deletion, almost completely abolished αII-spectrin detection (Fig. 5a). In the βII KO condition, the residual αII-spectrin labelling exhibited altered spatial organization characterized by a strong decreased in periodicity (i.e., reduced autocorrelation amplitude) and an impaired spacing as compared to control axons (Fig. 5a).

Together, these findings demonstrate that βII- and βIII-spectrins function as redundant but additive components of MPS organization in dendritic shafts and spines and indicate that the MPS is a structurally robust lattice capable of maintaining its integrity even when spectrin tetramer content is reduced.

### Distinct binding interactions govern βII- and βIII-spectrin dynamics and structural organization in the MPS lattice

Through evolution, duplicated genes can be preserved through subfunctionalization^44^, whereby paralogues acquire localization-, or stage-specific expression patterns, or through neofunctionalization^45^, in which one copy retains the ancestral function while the other evolves new functional properties. The coexistence of βII- and βIII-spectrins within the same subcellular compartments and at the same developmental stage argues against simple subfunctionalization and instead favors neofunctionalization. The robustness of MPS organization observed in our KO experiments further suggests that functional specialization of these paralogues may arise not from distinct structural roles in lattice assembly, but from their differential regulation within the MPS.

To test this hypothesis, we generated GFP-tagged βII- and βIII- spectrin constructs (GFP-βx WT) and mutant variants targeting interaction domains for specific binding partners. These included point mutants lacking ankyrin binding (βx Ank*), point mutants deficient in phosphoinositide interaction (βx PIP*), and deletion mutants lacking the actin-binding domains (βx dABD) (Fig. 6a). To avoid overexpression artifacts, we implemented a “re-expression” system by depleting endogenous βx-spectrin using CRISPR–Cas9 while co-expressing gRNA insensitive βx-spectrin constructs and their mutant variants at near-endogenous levels (Fig. S6).

**Figure 6.**
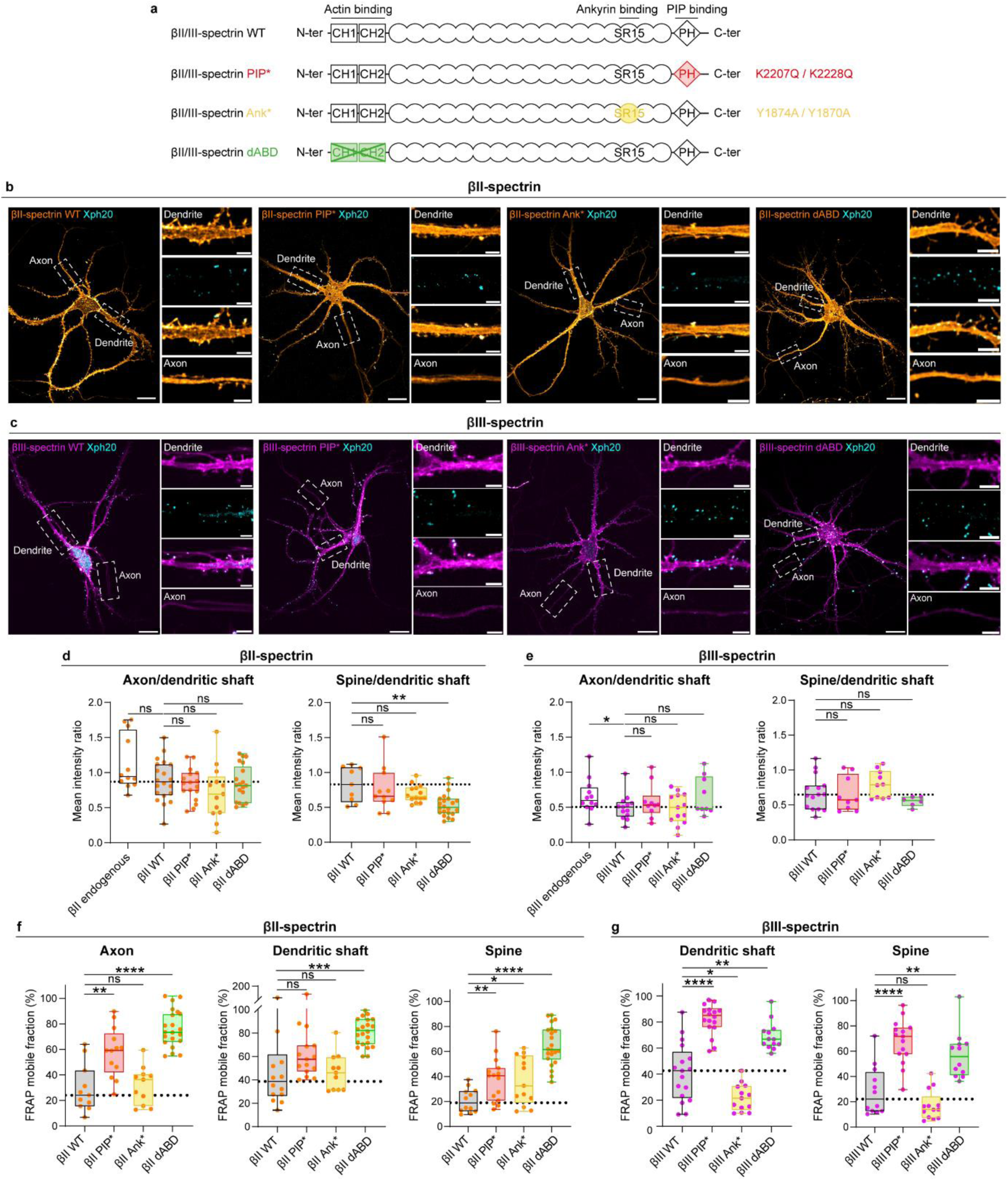
The binding partners of β-spectrins differently regulate their mobility. **a,** Scheme of βII- and βIII-spectrins highlighting the mutations disrupting interaction with phosphoinositides, ankyrins or actin. **b, c,** Representative spinning disk images of neurons re-expressing βII- (a) or βIII-spectrins (b) WT and mutant all tagged with eGFP, and expressing an intrabody labelling PSD95^76^. Scale bars in top images, 20 µm. Scale bars in bottom images, 1 µm. **d, e,** Ratios of the mean fluorescence intensity measured in axons, dendrites and spines in endogenous and re-expression conditions. n between 9 and 22 neurons obtained from 3 to 5 independent experiments for βII-spectrin, n between 7 and 16 neurons obtained from 3 to 4 independent experiments for βIII-spectrin. **f, g,** Mobile fractions of βII/III-spectrins measured 180 seconds after photobleaching. Each box represents the median with percentile. n between 12 and 24 regions of interest per neurite obtained from 3 to 5 independent experiments for βII-spectrin, n between 13 and 18 regions of interest per neurite obtained from 3 to 4 independent experiments for βIII-spectrin. Each box represents the median with percentile. Mann-Whitney tests, ns: non-significant, *: p < 0.05, **: p < 0.01, ***: p < 0.001, ****: p < 0.0001.

We first examined the effects of these mutations on βx-spectrin localization across neuronal compartments (Fig. 6). Axon/dendrite and spine/dendrite intensity ratios for the different mutants were largely comparable to their respective wild-type (WT) constructs, except for the βII-spectrin dABD mutant, which showed reduced accumulation in spines (Fig. 6d, e). At this resolution, the steady-state polarity of βII- and βIII-spectrins was minimally affected by the mutations (Fig. 6a–e).

We next assessed their dynamic interaction with the MPS lattice, analyzing their diffusion by fluorescence recovery after photobleaching (FRAP) (Fig. 6f, g and Fig.S7). Both βII- and βIII-WT displayed similar diffusion behavior, characterized by strong immobilization (i.e. small FRAP mobile fraction) in spines and dendritic shafts, with βII-spectrin showing comparable stabilization in axons, consistent with proper integration into the MPS lattice (Fig. 6f, g and Fig.S7). Actin binding proved essential for both paralogues, as the dABD mutants exhibited marked increases in mobility across all compartments (Fig. 6f, g). In contrast, βII- and βIII-spectrins differed in the relative contributions of ankyrin and phosphoinositide interactions to their stabilization. βII-spectrin stabilization was primarily governed by actin binding, with βII-Ank* and βII-PIP* mutants displaying intermediate phenotypes (Fig. 6f). βIII-spectrin, however, showed a pronounced dependence on phosphoinositide binding: the βIII-PIP* mutant exhibited markedly increased mobility compared with βIII-WT, exceeding that of the βIII-dABD mutant. Notably, the βIII-Ank* mutant showed the opposite behavior, displaying reduced mobility relative to WT, suggesting that, in the case of βIII-spectrin, ankyrin interaction impairs stabilization in dendritic shafts and spines (Fig. 6g).

To determine whether these differential interactions also influenced the nanoscale organization of β-spectrins within the MPS lattice, we examined the structural consequences of these mutations using super-resolution microscopy. *d*STORM imaging of βII- and βIII-spectrin mutants supported the conclusions drawn from the FRAP analysis. The periodic organization of both βII- and βIII-dABD mutants was disrupted across all compartments, as indicated by the combined significantly reduced autocorrelation amplitudes, in axons, dendritic shafts, and, for βIII-dABD, also in spine necks, as well as by the increased lattice spacing (in axons, dendritic shaft, and spines for βII-dABD, and in dendritic shafts for βIII-dABD) (Fig. 7). The nanoscale organization of βII-Ank* and βII-PIP* mutants was altered only in axons (Fig. 7a, c, e). For βIII-spectrin, mutant behaviors were consistent across dendritic shafts and spines. In both compartments, the βIII-PIP* mutant showed decreased autocorrelation amplitudes (statistically significant in dendritic shafts) and a pronounced increase in lattice spacing (statistically significant in dendritic shaft and spines) (Fig. 7b, d, f). Conversely, the βIII-Ank* mutant displayed increased autocorrelation amplitude (statistically significant in dendritic shafts) and reduced spacing (statistically significant in spines), consistent with its enhanced immobilization observed by FRAP in these two compartments (Figs. 6g and 7b, d).

**Figure 7.**
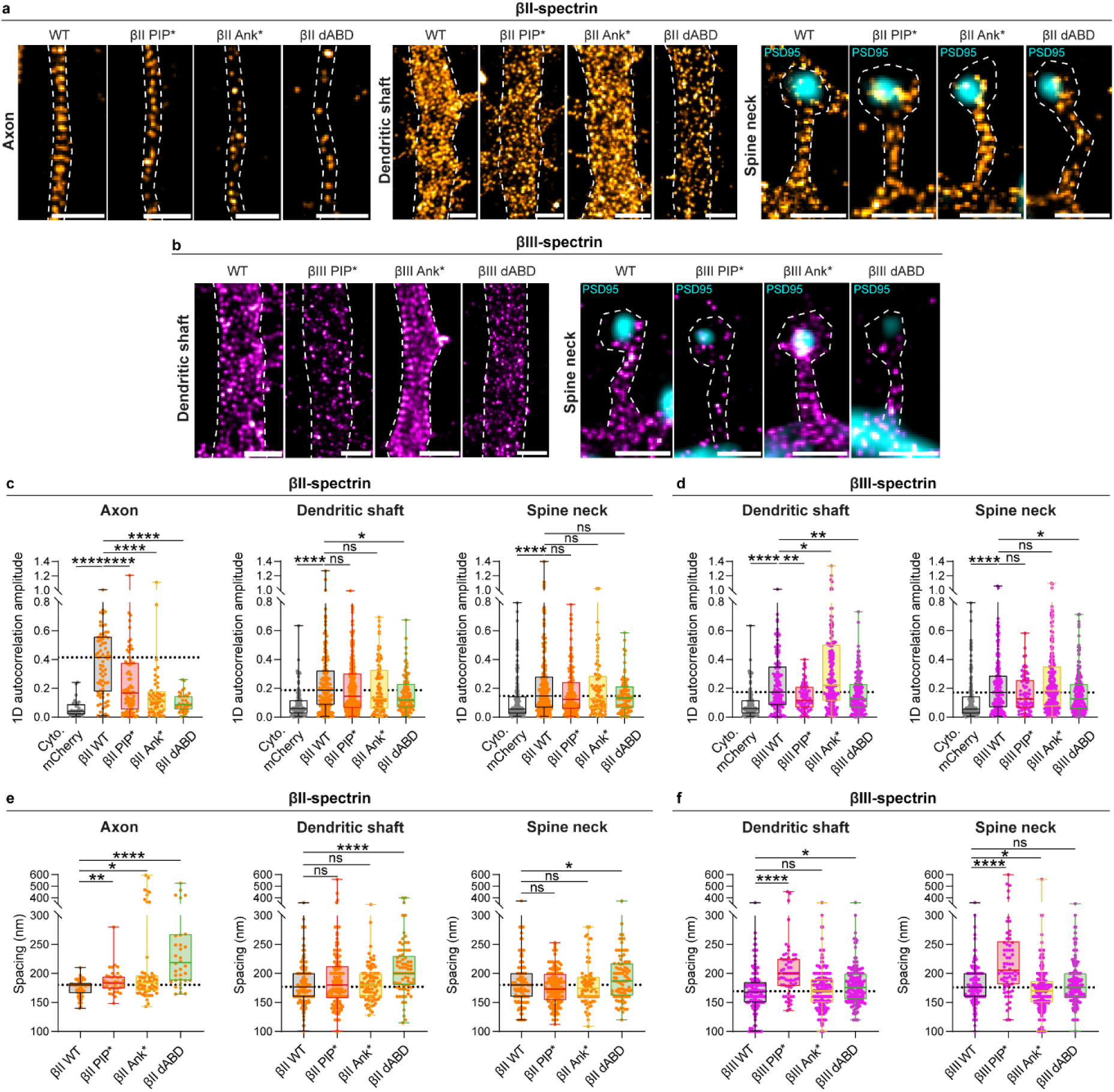
The binding partners of β-spectrins are differently involved in their periodic organization. **a, b** Representative *d*STORM images of wild-type and mutant βII-spectrins (a) and wild-type and mutant βIII-spectrins (b) imaged in re-expression condition in the indicated neuronal compartments. The non-periodic controls are the same as in figure 2. PSD95 was imaged using epifluorescence microscopy. Scale bars, 1 µm. **c, d** βII-spectrin (a) and βIII-spectrins (b) 1D autocorrelation amplitudes measured in the indicated compartments. n between 31 and 162 regions of interest per compartment obtained from 3 to 5 independent experiments for βII-spectrin. n between 73 and 213 regions of interest per compartment obtained from 3 to 4 independent experiments βIII-spectrins. Each box represents the median with percentile. Mann-Whitney tests, ns: non-significant, *: p < 0.05, **: p < 0.01, ****: p < 0.0001. **e, f,** Spacing between βII-spectrin (e) or βIII-spectrin (f) epitopes obtained from the same datasets as in (c) and (d).

Together, these results reveal both shared and distinct mechanisms governing βII- and βIII-spectrin organization in neurons. Actin binding is essential for the structural stabilization of both paralogues, whereas βIII-spectrin shows an additional and prominent layer of regulation through phosphoinositide interaction, and an opposing contribution of ankyrin binding. This differential reliance on specific binding partners suggests that βII- and βIII-spectrins contribute unequally to the regulatory and adaptive properties of the MPS lattice.

## Discussion

Our study reveals that the dendritic MPS is built from a dual β-spectrin system in which βII-and βIII-spectrin coexist and are intermixed within the MPS lattice (Fig. 8). Using advanced super-resolution imaging, our data suggest that the two paralogues assemble into both homotypic and heterotypic tetramers arranged with radial periodicity (Fig. 8). Despite this close spatial integration, selective depletion demonstrated that each paralogue is individually dispensable for maintaining MPS architecture, whereas loss of both led to collapse of lattice organization, establishing βII- and βIII-spectrin as redundant structural units. Importantly, preservation of MPS periodicity following single-paralogue depletion could not be attributed to compensatory protein upregulation, as neither β-spectrin paralogue increased upon loss of the other and αII-spectrin levels scaled with total β-spectrin availability. Thus, MPS persistence upon single β-spectrin depletion reflects genuine structural redundancy rather than molecular compensation. Finally, targeted mutagenesis uncovered paralogue-specific regulatory logic governing MPS integration: actin binding is essential for both β-spectrins, whereas βIII-spectrin uniquely depends on phosphoinositide interactions and is negatively regulated by ankyrin binding (Fig. 8). Together, these results identify a cooperative yet differentially controlled β-spectrin network that confers both robustness and regulatory flexibility to the dendritic MPS.

**Figure 8.**
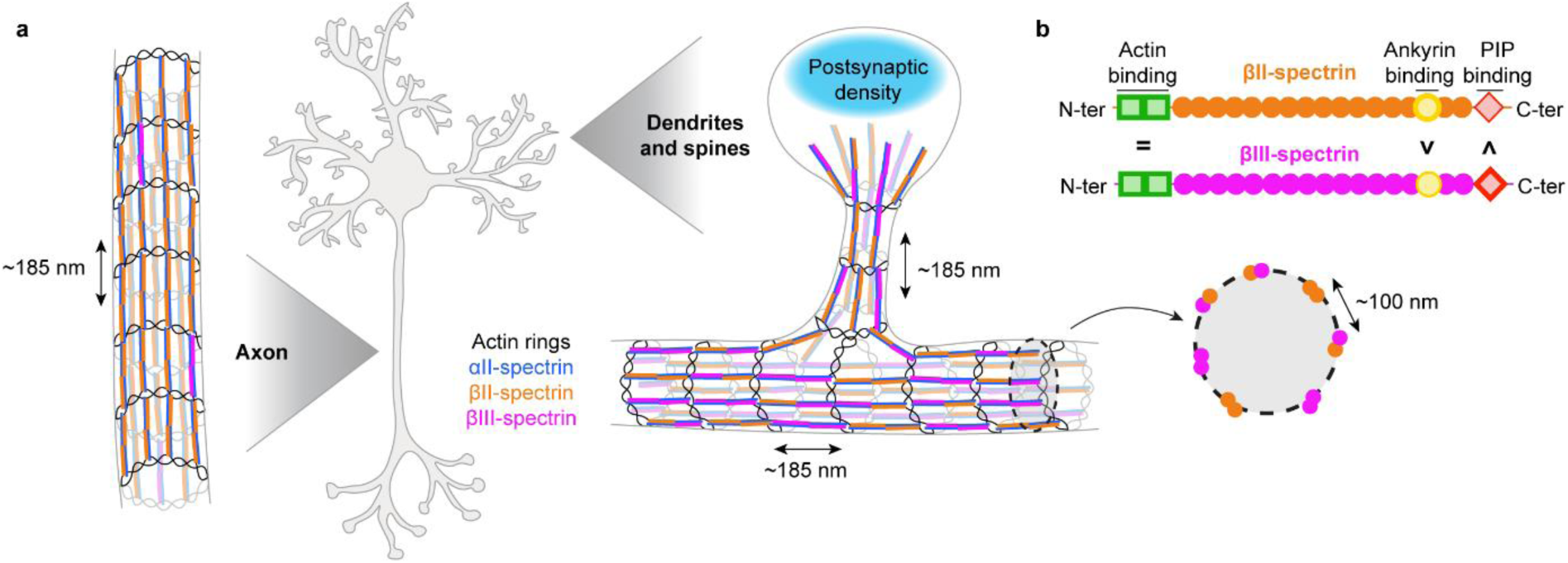
βII- and βIII-spectrins nano-organization in neurites and sensitivities to their binding partners. **a,** Schematic of the periodic actin-spectrin cytoskeleton in axons (βII-spectrin mostly), dendrites and dendritic spines (random βII and βIII spectrins distribution with the presence of sparse βII/βIII heterotypic tetramers). Spectrin signals are excluded from post-synaptic densities. **b,** Schematic of βII- and βIII-spectrins sensitivity to their binding partners.

Although the axonal MPS is well established as a strictly βII-spectrin-based lattice, much less was known about how β-spectrin paralogues contribute to MPS architecture in dendritic shafts and spines. Previous studies reported that both βII- and βIII-spectrin are present in dendrites^7,38^, but the extent of their nanoscale co-assembly remained unclear. Han and colleagues proposed that βIII-spectrin organization in dendritic shafts depends on βII-spectrin expression^7^, in contrast to our observations. This discrepancy likely reflects differences in experimental design. In their study, βII-spectrin was deleted at 3 DIV, when the dendritic MPS has not yet formed, whereas in our experiments deletion was induced later, at 8 DIV. These results suggest that βII-spectrin may contribute to the initial formation of the dendritic MPS but is not required for its maintenance once the lattice is established. Another important difference lies in the mode of genetic manipulation: Han and colleagues used viral infection^7^, resulting in βII-spectrin deletion in a large proportion of neurons, making it difficult to distinguish direct dendritic effects from indirect consequences of impaired axonal growth and altered synaptic connectivity in neighboring infected cells. By contrast, our sparse transfection strategy allowed manipulation of individual neurons within otherwise intact networks, enabling a more direct assessment of the cell-autonomous requirement of βII-spectrin for dendritic MPS organization. Even less was known about the organization of the MPS in dendritic spines. Periodic βII-spectrin organization had been reported in only a subset of spine necks (approximately 25–50%)^3,8^, often without extensive quantitative analysis. In contrast, nanoscale investigation of βIII-spectrin using platinum-replica electron microscopy didn’t detect periodic organization^38^. This absence of detectable periodicity likely reflects methodological limitations, as the same approach also failed to reveal periodic organization at the axon initial segment, where a canonical MPS is known to exist^46,47^. Consequently, it remained unclear whether dendritic spines contain a conventional actin-spectrin lattice, whether βIII-spectrin participates in its assembly, or whether dendritic MPS organization fundamentally differs from that of axons.

Together, these observations left the molecular composition and organization of the dendritic MPS unresolved. Our data address these ambiguities by demonstrating that both βII- and βIII-spectrin form periodic structures in dendrites and spine necks and are interleaved within the same lattice. These findings establish that the dendritic MPS is homogeneously dual in composition and structurally distinct from its axonal counterpart. In addition, our results suggest the coexistence of both homotypic and mixed-composition β-spectrin tetramers. However, unambiguous demonstration and quantitative assessment of heterotypic βII/βIII-spectrin tetramers will require future biochemical or structural analyses.

Another β-spectrin paralogue, the erythrocyte βI-spectrin paralogue, has been described in neurons, where it was found predominantly at the postsynaptic density and showed little to no co-localization with αII-spectrin in dendritic spines^48^. Immunoprecipitation experiments further suggested that βI-spectrin could exist predominantly in a monomeric form in this compartment^48^. Consistent with this view, we find that αII-spectrin signal in dendritic spines is completely lost upon combined depletion of βII- and βIII-spectrins (Fig. 5), arguing that βI-spectrin is not incorporated into αII-containing spectrin tetramers in these compartments and may function in a distinct, tetramer-independent role in spines.

Recent live super-resolution imaging has revised the view of the axonal MPS as a static structure, revealing constitutive cycles of local disassembly and reassembly driven by calcium signaling^49^. These dynamics were reduced by inhibition of action potentials, indicating tight coupling between neuronal activity and MPS remodeling in axons^49^. Previous work using high extracellular potassium as a depolarization paradigm showed that dendritic, but not axonal, actin rings underwent large-scale reorganization into longitudinal actin fibers in a calcium-dependent manner^50^. Together, these observations indicate that while both axonal and dendritic MPS structures are dynamically regulated, dendrites may be more prone to activity-driven remodeling than axons.

The reasons underlying the resistance of the axonal lattice to extracellular potassium stimulation remain to be determined. Differential MPS composition could contribute, or alternatively, calcium elevation in axons may be attenuated relative to other neuronal compartments. Dendrites and dendritic spines experience strong local calcium transients that are highly heterogeneous in space, amplitude, and duration, reflecting the diversity of intrinsic neuronal excitability and synaptic activity^51^. Structural redundancy between β-spectrin paralogues could ensure lattice persistence and prevent catastrophic collapse during frequent calcium elevations, while paralogue-specific regulatory features may expand the range of possible remodeling responses to activity-dependent signaling pathways. Consistent with this model, βII- and βIII-spectrins are not stabilized identically in dendritic shafts and spines. Whereas actin binding is required for both isoforms, βIII-spectrin depends strongly on phosphoinositide interactions and is destabilized by ankyrin binding. Given the dynamic and spatially heterogeneous regulation of phosphoinositide composition in neurons during synaptic plasticity^52–54^, selective destabilization of βIII-containing assemblies could, for example, enable localized and reversible weakening of the lattice while βII-spectrin maintain the overall structural integrity.

Localized remodeling of the dendritic MPS is likely to have direct consequences for membrane trafficking and diffusion. The actin-spectrin lattice acts as a physical barrier to endocytosis^20,55,56^, restricting clathrin-coated pit formation to MPS-free membrane domains. In the confined geometry of dendritic spines, differential regulation of β-spectrin paralogues may therefore contribute to coupling between endocytosis and synaptic activity^57–59^. In addition, a spectrin-based cytoskeleton closely apposed to the plasma membrane has been shown to restrict lateral diffusion of transmembrane proteins and lipids in erythrocytes and axons^60–65^. By analogy, the dendritic MPS could modulate the mobility of synaptic receptors such as AMPA receptors in a tunable manner depending on the strength of interactions with specific phosphoinositides in the inner leaflet of the membrane, thereby influencing synaptic transmission and plasticity^66–69^. Although we do not directly assess activity-dependent membrane trafficking or diffusion in this study, our findings suggest that spectrin-dependent regulation may fine-tune endocytosis and protein mobility, with potential consequences for synaptic function.

From an evolutionary perspective, the mammal-specific emergence of βIII-spectrin^23,24^ can be interpreted in the context of a broader shift in how adult vertebrate nervous systems handle plasticity and repair. In many non-mammalian species, plasticity and recovery from damage can involve changes at the level of circuit composition, including the addition or replacement of neurons^70^. In mammals, by contrast, neuronal replacement and regenerative repair in the central nervous system are strongly constrained, and plasticity relies predominantly on the refinement of pre-existing circuits through synaptic and subcellular remodeling within long-lived neurons^70,71^. This strategy places strong demands on cytoskeletal systems that must preserve neuronal architecture over long timescales while remaining compatible with ongoing activity-dependent remodeling. Within this framework, the periodic actin-spectrin scaffold built from βII- and βIII-spectrins is well suited to such a role. Our results are consistent with a model in which βIII-spectrin contributes a specialization of the dendritic MPS, potentially providing an additional layer of regulation within a structurally robust dendritic MPS, helping to reconcile long-term stability with continued synaptic remodeling under limited neurogenic capacity in mammals.

Pathogenic variants in spectrin genes are associated with neurodevelopmental disorders and, although these variants are classified as rare, their identification in individuals with neurological conditions is rapidly increasing^72^. Variants in βII-spectrin and βIII-spectrin have been genetically and functionally linked to neurological disorders whose clinical manifestations include intellectual disability, developmental delay, seizures, movement disorders, and behavioral abnormalities^72^. Several disease-associated βII- and βIII-spectrin variants cluster in the calponin homology domains and dysregulate binding to F-actin. Notably, pathogenic variants affecting the pleckstrin homology domain have so far been described only for βIII-spectrin^72,73^, further supporting a paralogue-specific role for this domain in βIII-spectrin function.

Altogether, our results identify the dendritic MPS as a dual β-spectrin scaffold that combines structural redundancy with paralogue-specific regulatory features. This organization provides a mechanistic framework through which dendrites may preserve long-term structural integrity while remaining permissive to localized, activity-dependent remodeling. By linking β-spectrin composition, molecular interactions, and nanoscale organization, our findings open new avenues to investigate how neuronal activity shapes the membrane-associated cytoskeleton and how alterations in this system contribute to synaptic dysfunction and neurodevelopmental disease.

## Material and Methods

### Animals

All procedures involving animals were done in accordance with the European Union Directives (2010/63/EU) and the University of Bordeaux Ethical Committee and French Research Ministry. Sprague-Dawley rats (Janvier Labs) from both sexes were used at E18.

### Plasmids

The GFP- βII-spectrin plasmid was a gift from Van Bennett (Duke University, USA). Myc-beta3-spectrin was a gift from Adam Avery (Oakland University, USA). Mutations abolishing the interaction with phosphoinositides and ankyrin were based on previous characterization of SPTBN1^74,75^ (βII-spectrin) and translated to the corresponding amino acid in the SPTBN2: SPTBN1-K2207Q and SPTBN2-K2228Q for the PH domain, SPTBN1-Y1874S and SPTBN2 Y1870A for the ankyrin binding domain. Mutagenesis was used to generate SPTBN1 mutants and nucleotides synthesis to replace the SPTBN2 (βIII-spectrin) wild type sequence by a mutated one (Table 1). Both CH1/2 actin binding domains were removed from the SPTBN1/2 sequence to abolish the interaction with actin by nucleotides synthesis (Table 1). mCherry sequence was added to the N-terminal part of all spectrins ORF to generate fusion constructs in a standard CMV promoter mammalian expression vector.

**Table 1.**
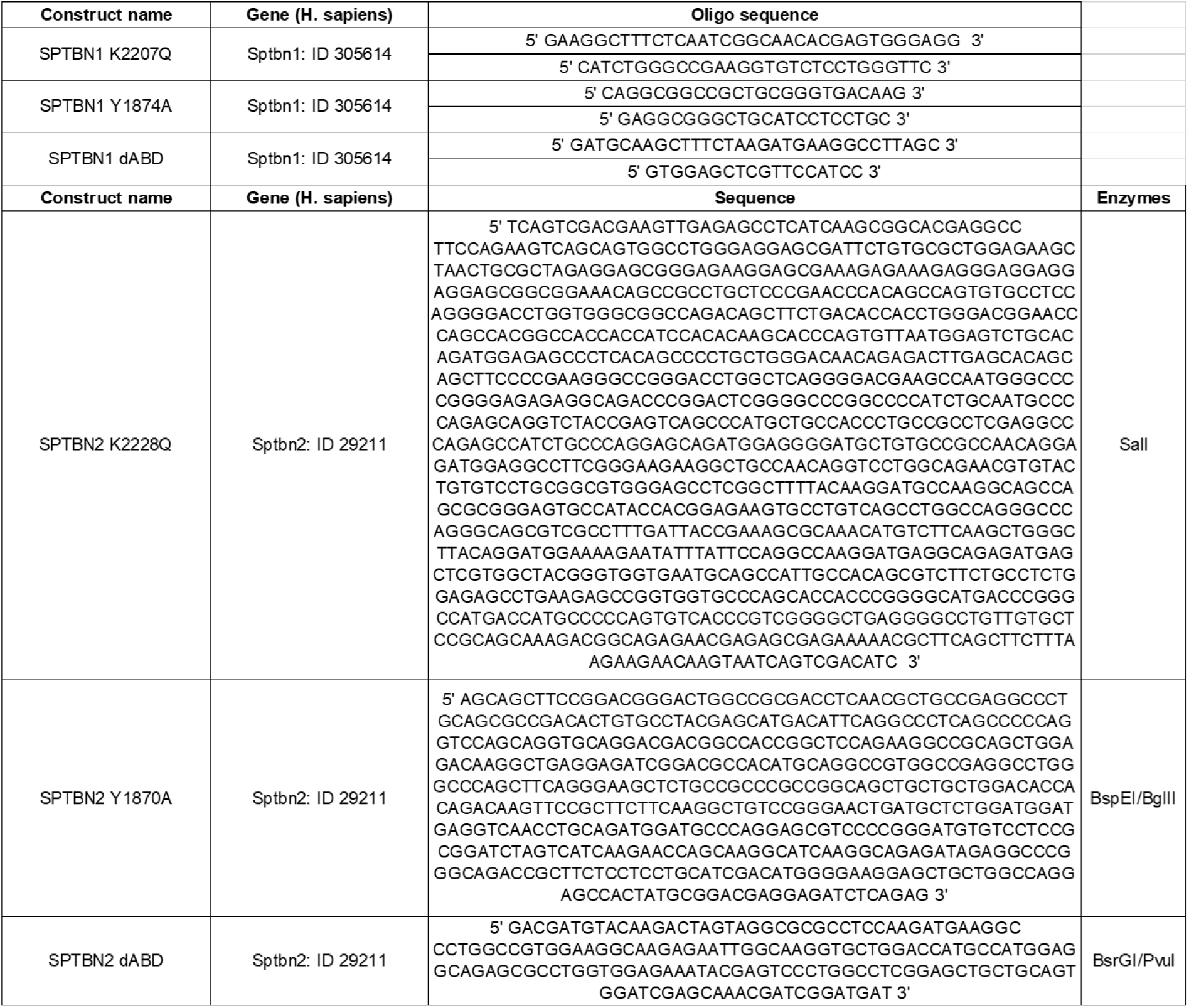

For CRISPR-Cas9-mediated gene inactivation, the choice of guide RNAs (gRNAs) was done using MIT (http://crispr.mit.edu) and Chochop (https://chopchop.cbu.uib.no/). To construct gRNA expression vectors, a pair of complementary 20 bp oligos for each gRNA sequence was annealed and cloned into all in one vector pSpCas9(BB)-2A-GFP (PX458) (Addgene plasmid # 48138; http://n2t.net/addgene:48138 ; RRID:Addgene_48138), using BbsI restriction site (Table 2). All the constructs were confirmed by sequencing. The control, non-targeting guide RNA (gRNA CTL) is a gift from CRISPR’EDIT (University of Bordeaux), obtained from a Gecko screen. An 2A-EGFP or tdTomato sequence was inserted after the S. pyogenes Cas9 (SpCas9) sequence as a transfection reporter.

**Table 2.**
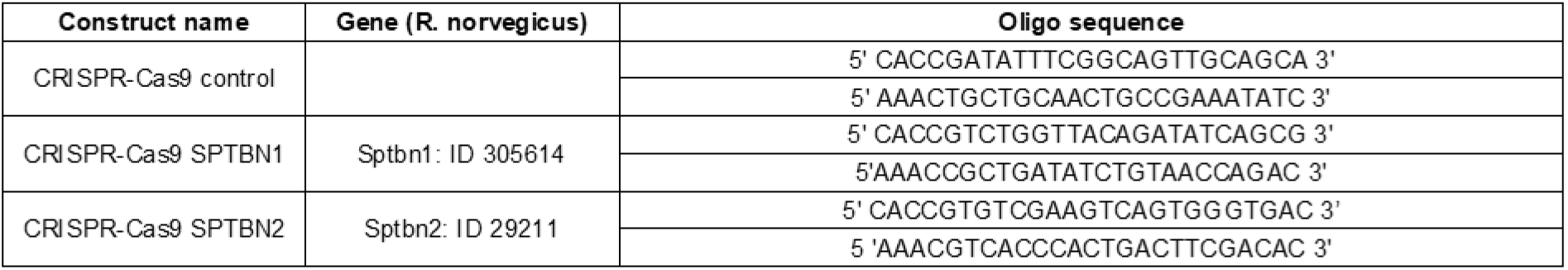

The CAAX-GFP plasmid was a gift from René-Marc Mège (Institut Jacques Monod, Paris). The Xph20-mRuby2 plasmid was a gift from Matthieu Sainlos^76^ (Interdisciplinary Institute for Neuroscience, Bordeaux, France).

### Antibodies

The following primary antibodies were used: βII-spectrin (BD Biosciences, 612 563, mouse IgG1 monoclonal, 1/500), βIII-spectrin (Novus Biotechnologies, NB110 58346, rabbit polyclonal, 1/500 or Santa Cruz, sc515737, monoclonal, 1/100), αII-spectrin (Biolegend, 803206, mouse IgG2b monoclonal, 1/1000), α-Adducin (Abcam, 51130, rabbit IgG polyclonal, 1/200), GFP (Avès, GFP-1020, chicken monoclonal, 1/2000), DsRed (Takara, 632496, rabbit polyclonal, 1/2000), PSD-95 (ThermoFisher, MA1-046, mouse IgG1 monoclonal, 1/500), Homer 1C (Synaptic Systems, 160004, guinea pig polyclonal, 1/500).

The following secondary antibodies were used (all of them at 1:500): anti-mouse AlexaFluor 647 (ThermoFisher A21235, goat), anti-rabbit AlexaFluo 647 (ThermoFisher, A21244, goat), anti-mouse CF680 (Sigma, SAB4600394), anti-rabbit CF680 (Sigma, SAB4600362), anti-mouse AlexaFluor 532 (ThermoFisher, A11002, goat), anti-guinea pig Dylight 405 (Jackson, 103-475-003, goat), anti-mouse IgG2a AlexaFluor 350 (ThermoFisher, A21130, goat), anti-chicken AlexaFluor 488 (ThermoFisher, A11039, goat), anti-rabbit AlexaFluor 568 (ThermoFisher, A11011, goat).

### Primary dissociated hippocampal culture and transfection

Dissociated hippocampal neurons from embryonic day 18 (E18) Sprague-Dawley rats of both sexes were prepared as previously described^77^. Briefly, dissociated neurons were plated at a density of 250,000 cells per 60 mm dish on 1 mg mL^−1^ PLL pre-coated 1.5H, ⌀ 18 mm coverslips (Marienfeld Superior, #0117580). Neuronal cultures were maintained in Neurobasal™ Plus Medium (Thermo Fisher Scientific) supplemented with 0.5 mM GlutaMAX (Thermo Fisher Scientific) and 1X B-27™ Plus Supplement (ThermoFisher Scientific). Cultures were kept at 37 °C under 5% CO_2_ up to 21 days. Neurons were transfected with the cDNA using Ca^2+^ method at 8–11 days in vitro.

### Immunocytochemistry

Primary hippocampal neurons aged from 10 to 21 DIV were fixed with 4% PFA with sucrose for 10 min at RT, before quenching the free aldehyde groups for 5 min (50 mM NH_4_Cl in 1X PBS). After 3 washings in 1X PBS, neurons were permeabilized (0.2% Triton X100, 1% BSA, 0.5% fish gelatin in 1X PBS) before incubating for 1 hour in a blocking buffer (1% BSA, 0.5% fish gelatin in 1X PBS). The cells were then incubated with primary antibodies for 2 hours at RT, washed 3 times in 1X PBS, and incubated with secondary antibodies for 45 min at RT. Coverslips were mounted on glass slides with Fluoromount.

### Wide-field microscopy

Neurons were imaged with a Leica DM5000 up-right widefield microscope (Leica Microsystems, Wetzlar, Germany) using the HCX PL APO 63X oil NA 1.40 objective and a Flash4.0 V2 camera (Hamamatsu Photonics, Massy, France). Fluorescence excitation was generated by a LED SOLA Light (Lumencor, Beaverton, USA) and the z stacks were done with a galvanometric stage (Leica Microsystems, Wetzlar, Germany). Measurements of fluorescence intensity in somas, axons, dendrites and spines were performed manually on a single plane using Fiji (https://imagej.net/Fiji).

### Confocal microscopy

Neurons were imaged using a Leica SP8 WLL2 confocal microscope on an inverted stand DMI6000 (Leica Microsystems, Mannheim, Germany) using objective HCX Plan Apo CS2 63X oil NA 1.40. The confocal microscope was equipped with a pulsed white light laser 2 (WLL2) with freely tuneable excitation from 470 to 670 nm (1 nm steps) and 405 nm laser diode. The scanning was done using a conventional scanner (10 Hz to 1800 Hz) Measurements of fluorescence intensity in dendrites and spines were performed using Fiji (https://imagej.net/Fiji).

### Fluorescence Recovery After Photobleaching (FRAP) imaging

Neurons were imaged at 14-17 DIV for FRAP imaging. FRAP imaging was performed on a spinning disk microscope Leica DMI8 (Leica Microsystems, Wetzlar, Germany) equipped with a confocal Scanner Unit CSU-W1 T2 (Yokogawa Electric Corporation, Tokyo, Japan) with a 63X oil objective (Leica, HC PL APO CS2, NA 1.4). The LASER diodes used were 491 nm (400 mW), 561 nm (400 mW) and 642 nm (100 mW). Images were acquired using a sCMOS Prime 95B camera (Photometrics, Tucson, USA). Photo-bleaching was done with an iLas² scanner system (Gataca Systems, Massy, France). The 37°C atmosphere was created with an incubator box and an air heating system (PeCon GmbH, Germany). Experiments were performed in Tyrode buffer (D+ Glucose 10 mM, HEPES 10 mM, NaCl 120mM, KCl 3.5 mM, MgCl_2_:6H20 2mM, CaCl_2_:2H20 2mM) at 37°C. eGFP and mRuby2 images of neurons were acquired before FRAP imaging to identify the regions of interest (ROIs) for bleaching. For photobleaching we used 5 ms pulse of 491 nm laser light sufficient to reduce fluorescence by at least 50%. Images were taken every 0.2 seconds for 10 seconds, then every 2 seconds for 60 seconds and finally every 30 seconds for 5 min. FRAP experiments were analyzed using an in-house-developed macro to the ImageJ freeware (http://imagej.nih.gov/ij/). The source code is freely available from GitHub (https://github.com/fabricecordelieres/IJ-Macro_FRAP-MM), accompanied by a documentation and an example dataset. Briefly, as part of the macro, the FRAP region for each spine was imported from the MetaMorph software to ImageJ’s ROI Manager using the “MetaMorph Companion” plugin (https://github.com/fabricecordelieres/IJ-Plugin_Metamorph-Companion). From the data extracted by the macro, average intensity within the ROI was collected for each timepoint (*F*_t_), at first timepoint (pre-bleach, *F*_pb_), and immediately after the bleaching (*F*_0_). Simple normalization was performed as follows: FRAP_t_ = *F*_t_ − *F*_0_/*F*_pb_ − *F*_0_. The mean spine FRAP curve of each cell was subsequently fitted to a monoexponential model using GraphPad Prism software. The mobile fraction corresponds to the plateau of the fitted curve.

### Immunocytochemistry for dSTORM and 3D MINFLUX imaging

After 21 DIV, primary neuronal cultures were fixed with PFA-sucrose (4%) for 10 minutes at RT and then washed 3 times with 1X PBS. PFA was quenched with NH_4_Cl 50 mM for 5 min and then washed 3 times with 1X PBS. After blocking and permeabilizing with blocking solution (0.1 % Triton 10X, 1% BSA, 0.5% fish gelatin in PBS 1X) for 1h, cells were incubated with primary antibodies overnight at 4°C and then washed with blocking solution three times for 5 min. Cells were then incubated with secondary antibodies for 45 min at RT and rinsed with blocking solution 3 times for 5 min and once with 1X PBX. The coverslips were then fixed a second time with PFA 2% for 10 minutes and then washed 3 times with 1X PBS. PFA was quenched with 50 mM NH_4_Cl for 5 min. Finally, the coverslips were rinsed 3 times with 1X PBS and kept at 4°C until imaging. For two-colour dSTORM and MINFLUX experiments, one β-spectrin was labelled by a secondary antibody conjugated to AlexaFluor647 and the second β-spectrin was labelled by a CF680-coupled secondary antibody. To localize dendritic spines a secondary antibody conjugated to AlexaFluor488 was used to label of Homer protein.

### direct STochastic Optical Reconstruction Microscopy (*d*STORM)

*d*STORM imaging was performed on a commercial inverted microscope (DMi8, Leica, Germany) equipped with a HC PL APO 100X oil immersion TIRF objective (1.47 NA, Leica), a Ilas² TIRF device, an Evolve EMCCD camera (Tedelyne Photometrics) and laser modules with following wavelength: 405, 488, 561, 642 nm (Coherent, USA). Experiments were performed as previously described^78,79^. Briefly *d*STORM acquisition was done in photoswitching buffer containing a reducing agent and an oxygen scavenger. Ensemble fluorescence of Alexa-647 was first converted into dark state using a 642 nm laser (Coherent) at 30 –50 kw/cm2 intensity, the laser power was then reduced to 7–15 kw/cm2. Acquisition was performed with an exposure time of 30ms and 20,000 frames were acquired. Multi-color fluorescent microspheres (Tetraspeck, Invitrogen) were used as fiducial markers to register long-term acquisitions and correct for lateral drifts. Super-resolved images were reconstructed with a 40 nm pixel size, using PALMTracer software, a MetaMorph (Molecular Devices, Sunnyvale, USA) add-on developed at the Interdisciplinary Institute of Neuroscience (IINS - UMR5297 -CNRS / University of Bordeaux), by Corey Butler(IINS -UMR5297 -CNRS / University of Bordeaux), Adel Mohamed Kechkar (Ecole Nationale Supérieure de Biotechnologie, Constantine, Algeria) and Jean-Baptiste Sibarita (IINS -UMR5297 -CNRS / University of Bordeaux). Briefly, Single molecule localization was achieved using wavelet segmentation, and then filtered out based on the quality of a 2D gaussian fit^80–82^.

### Two-color *d*STORM imaging

Two-color *d*STORM imaging was performed on a commercial inverted microscope (DMi8, Leica, Germany) equipped with an Abbelight system containing a SAFe360 module and an ASTER scanning module (Abbelight, Cachan, France). This system is combined with two sCMOS ORCA-Fusion BT cameras for the transmitted light and for the reflected light (Hamamatsu, Japan) and the following lasers: 405, 488, 532, 561 and 642nm. Acquisition was driven by Neo Live software. Low resolution images were acquired in the different channels before acquisition. For acquisition, exposure time was set up at 50 ms and 40 000 frames were registered to reconstruct the final super-resolved images. This setup allows performing two-color *d*STORM imaging using spectral demixing technique and hence avoid chromatic aberration that can be encountered on the setup described above^83^. For simultaneous dual protein observations, spectral demixing was used (SAFe360 module). Briefly spectrally close fluorophores (here AF647 and CF680) were illuminated by a single laser as previously described^84^. The emission fluorescence is divided into two beams by a dichroic mirror with a cut-off wavelength at 700 nm (Chroma T700lpxr-3). Each beam is directed to two separate cameras (transmitted and reflected pathways), two recordings are thus acquired that can be further demixed with ratiometric analysis to obtain two-color super-resolved images. Analysis, including drift correction and ratiometric measurements for spectral demixing were performed with Neo analysis software (Abbelight, Cachan, France).

### Autocorrelation, cross-correlation analysis and spacing

Autocorrelation and cross-correlation measurements were done using the “Correlate Profiles” script and protocol from C. Leterrier^47^. The script is freely available at https://github.com/cleterrier/Process_Profiles

The autocorrelation function evaluates how similar a signal is to a delayed version of itself, revealing periodicities or repeating patterns within the signal. In contrast, the cross-correlation function assesses the degree of similarity between two distinct signals, as a function of their relative shift, allowing the identification of spatial alignment between them.

Briefly, line regions were defined on identified spine necks and neighbouring dendrites on *d*STORM reconstructed images with a pixel size of 40nm. The used script allowed to extract the normalized autocorrelation curve for each intensity profile. For every autocorrelation curve, the first peak was identified using a home-made ImageJ macro, to extract the corresponding spacing value, here 188 nm measured for βII-spectrin in axons for instance. The autocorrelation amplitude which corresponds to the height of the first peak of the curve, was calculated by the difference of the autocorrelation value at 188 nm (position of the first peak) and the average value at 94 nm and 282 nm (first and second valleys).

Autocorrelation amplitude reports the global coherence of the periodic pattern, integrating the spatial organization over a certain length, while spacing is sensitive to local positional variability. Both parameters are therefore complementary measurements to check for MPS integrity.

### MINFLUX 3D microscopy

Samples were imaged on two Abberior Instruments MINFLUX microscopes.

Details of the system layout and components are described in Schmidt et al., 2021. Briefly, the microscopes were built on motorized inverted microscope bodies (IX83, Olympus) with a CoolLED pE-4000 (CoolLED) for epifluorescence illumination, and equipped with 405 nm, 485 nm, 561 nm, and 642 nm laser lines, as well as a 980 nm IR laser and xyz piezo (Piezoconcept) for the active sample stabilisation system. The reflection from the 980 nm laser were recorded by the dedicated stabilisation system and used to assess sub-nanometer shifts in the sample position. These deviations were corrected actively during the measurement with the xyz piezo stage. Both systems contain a 642 nm MINFLUX laser. Images were acquired with a 100x 1.45 NA UPLXAPO oil objective (Olympus) or 60x 1.42 NA UPLXAPO oil objective (Olympus). The ROIs were selected in the confocal mode.

The three dimensional spectral demixing MINFLUX images (3D-SD MINFLUX) were recorded with a pinhole diameter of 0.8 Airy Units and detected across two spectral windows from 650 nm – 685 nm (Cy5 near) and 685 - 720 nm (Cy5 far). As with two color dSTORM imaging, photons were received over two detectors, allowing for spectral demixing of the distinct populations on a ratiometric basis. MINFLUX laser power in the first iteration was measured at the periscope to be ∼0.2 mW.

Samples for spectral demixing MINFLUX were incubated with 150 nm gold beads (BBI Solutions) for 5 minutes before washing three times with PBS, with a final 5 minute incubation in PBS supplemented with 10 mM MgCl2. The coverslips were then mounted on a cavity slide (BMS Microscopes) in a STORM blinking buffer (pH 8.0) composed of 50 mM Tris-HCl, 10 mM NaCl, 10% Glucose (w:v), 64 µg/ml catalase (Sigma), 0,4 mg/ml glucose oxidase (Sigma) and 53-63 mM MEA (Sigma). Samples were sealed against oxygen with a two part silicone (eco-sil speed, picodent).

Imaging was performed with the default 3D imaging sequence supplied with the system. In brief, the sequence defines parameters used in the iterative localising process of MINFLUX, culminating in a final pair of lateral and axial iterations with a TCP (targeted coordinate pattern) diameter of 40 nm, and minimum photon thresholds of 100 and 50 photons, respectively.

For MINFLUX beamline monitoring (mbm), 7-9 gold beads were selected as reference markers and used to correct any system drift over the course of the measurement.

After data acquisition, data was filtered by the EFO (effective frequency at the offset, EFO < 80 - 110 kHz). Detections were assigned to the relevant fluorophore population by definition of threshold values for the DCR (detector channel ratio, calculated as: counts on detector 1/ counts on detectors 1 + 2) based on the histogram of the DCR values of the last MINFLUX iteration. Thresholds were defined between peaks in the histogram, with mixed populations between peaks excluded. MINFLUX images were exported as TIFF stacks with 5 nm voxel size before 3D rendering and analysis in Imaris.

## Statistical analysis

All results are derived from at least three independent experiments. For imaging data, analysis was performed blinded to the experimental condition. Statistical analysis and data plotting was performed with Prism 9.0.2 (GraphPad). Statistical test details are reported in the figures’ legends.

## Acknowledgments

We would like to thank Christophe Leterrier (INP UMR7051, Université Aix Marseille, France) for insightful discussions and feedback on the manuscript and Jean-Baptiste Sibarita (IINS UMR 5297, Université de Bordeaux, France) for the PALMtracer software.

The microscopy was done in the Bordeaux Imaging Center a service unit of the CNRS-INSERM and Bordeaux University, member of the national infrastructure France BioImaging supported by the French National Research Agency (ANR-10-INBS-04). The help of Fabrice Cordelières is acknowledged for his help on FRAP experiments analysis.

We thank the IINS Cell Biology Facility used in this study, supported by the GPR BRAIN_2030.

## Author contributions

Conceptualization: AB, MLJ, LS

Methodology: AB, MLJ, LS, MM, IJ, EG, SD, PB

Investigation: AB, MLJ, LS, MM, IJ, EG, SD, PB, NC, TC

Supervision: AB, MLJ

Writing, original draft: AB, LS, MLJ

Writing, review & editing: AB, LS, MLJ, MS, DC, EG, IJ, NC, SD, MM

## Data and materials availability

All data are available in the main text or the supplementary materials.

