## Supplementary figures for "βII and βIII spectrin paralogues define robustness and specialization of the neuronal membrane periodic skeleton"

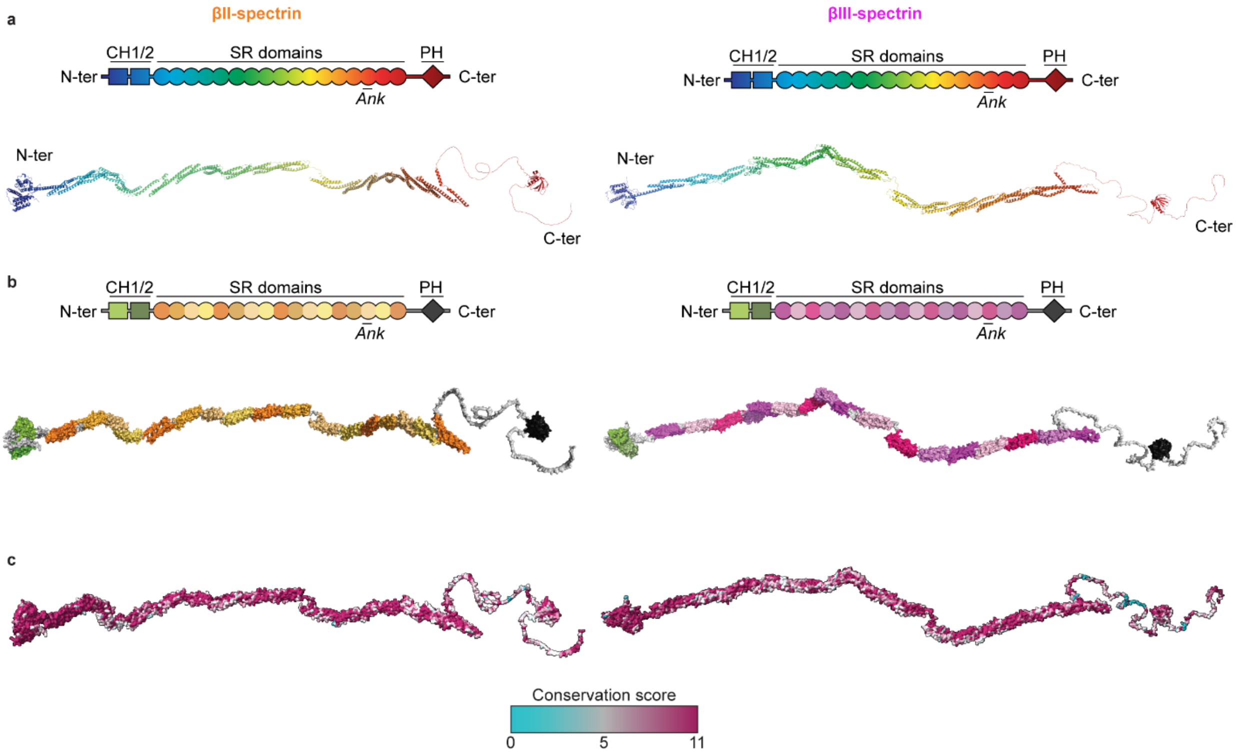


**Figure S1. Molecular architecture of βII and βIII spectrins and structural domains conservation. a,** Alphafold3 structure prediction of βII (Uniprot Q01082) and βIII (Uniprot O15020) spectrins. **b,** Main functional domains organization. **c,** Functional domains conservation score (βII versus βIII comparison) mapped by amino acid onto spectrin structures.

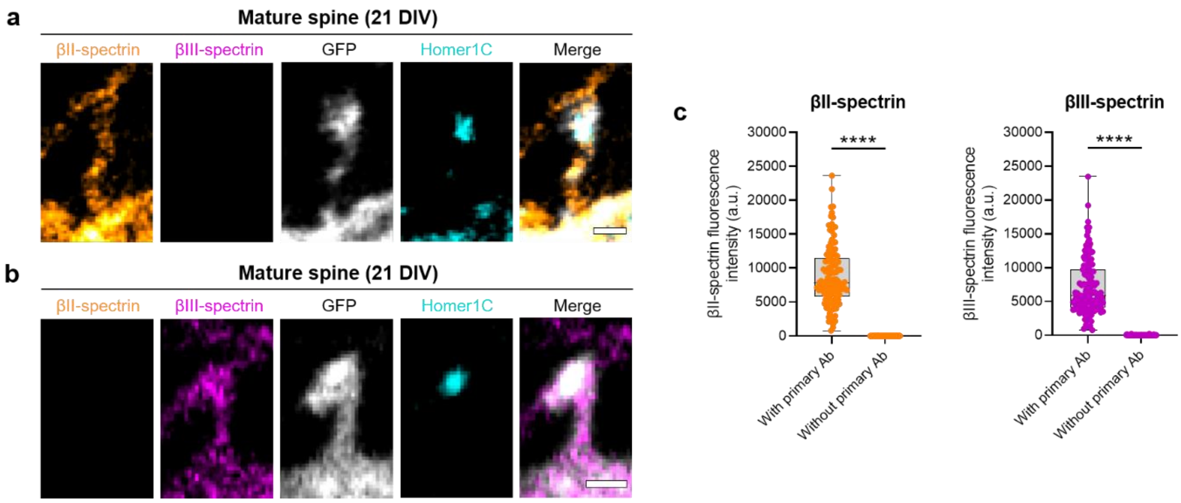


**Figure S2. Fluorescence bleed-through between βII- and βIII-spectrins is negligible. a, b,** Representative confocal images of βII-spectrin (orange), βIII-spectrin (magenta), CAAX-GFP (grey) and Homer1C (cyan) in mature spines at 21 DIV, immuno-labelling in the absence of the primary antibody for either βIII-spectrin (a) or βII-spectrin (b). Scale bars, 1 µm. **c,** Fluorescence intensity measurement of βII-spectrin or βIII-spectrin in spine necks at 21 DIV with or without the corresponding primary antibody. Each box shows the median ± percentile. n between 70 and 162 spine necks from 6 to 24 neurons examined over at least 2 independent experiments. Mann-Whitney test, ****: $p < 0.0001$.

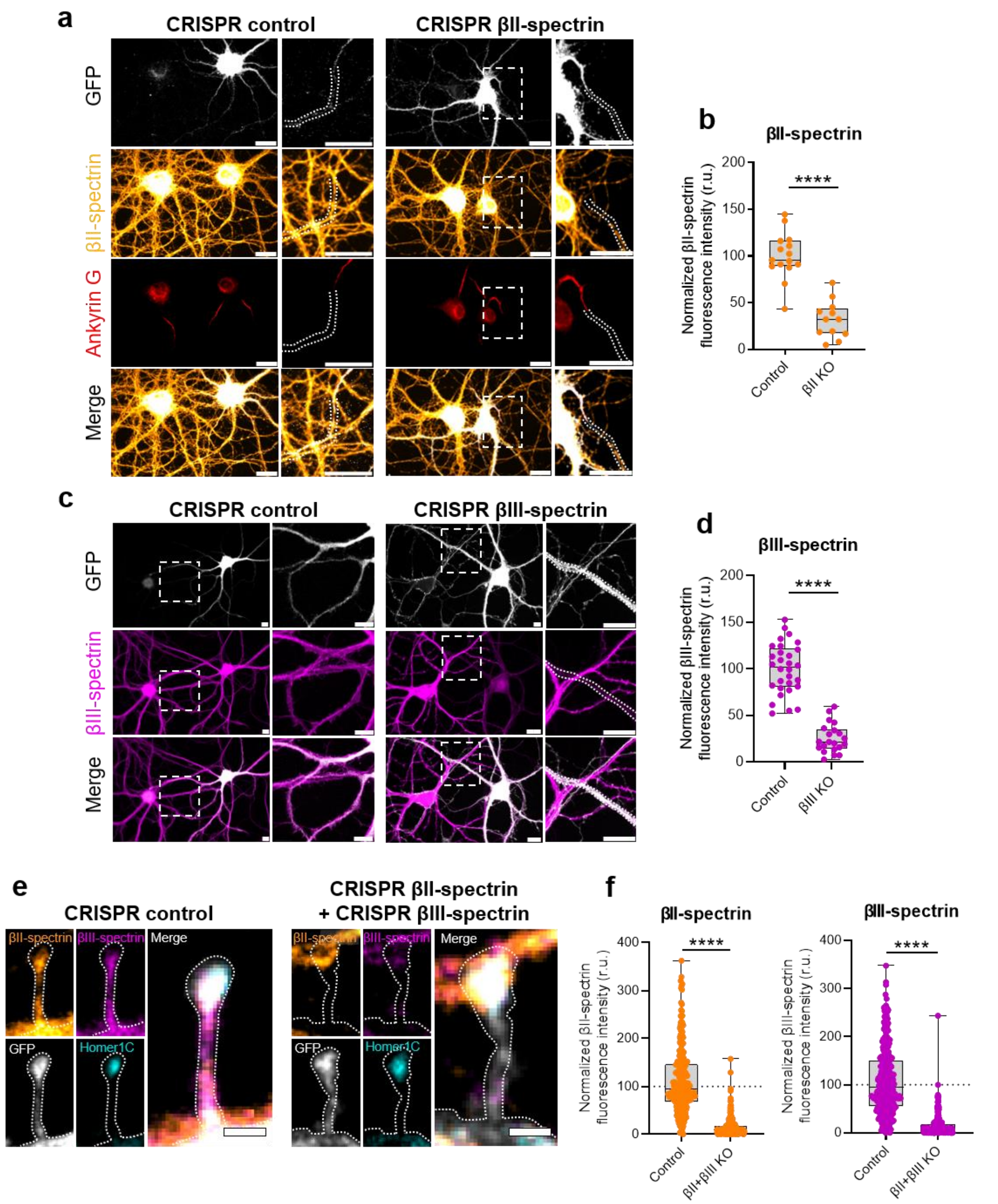


**Figure S3. CRISPR-Cas9 strategies targeting βII- or βIII-spectrin are efficient. a,** Representative wide-field images of a non-transfected neuron (GFP-negative) alongside a neuron transfected with the CRISPR-Cas9 system and a gRNA control (left) or targeting a βII-spectrin sequence (right). βII-spectrin fluorescence was assessed in isolated axonal regions following the ankyrin G signal, such as framed in bold white, to avoid signal contamination by neurites from neighbouring neurons. Scale bars, 10 µm. **b,** Quantification of βII-spectrin fluorescence intensity in axons of control versus βII KO neurons. Each box represents the median with percentile. n = 15 and 12 neurons from 3 independent experiments. Mann-Whitney test, ****: $p < 0.0001$. **c,** Representative wide-field images of a non-transfected neuron

(GFP-negative) alongside a neuron transfected with the CRISPR-Cas9 system and a gRNA control (left) or targeting a βIII-spectrin sequence (right). βIII-spectrin fluorescence was assessed in dendrites, such as framed in bold white. Scale bars, 10 µm. **d,** Quantification of βIII-spectrin fluorescence intensity in dendrites of control versus βIII KO neurons. n = 30 and 21 neurons from 3 independent experiments. Mann-Whitney test, ****: $p < 0.0001$. **e,** Representative confocal images of dendritic spines from a neuron transfected with the CRISPR-Cas9 system and a gRNA control (left) or gRNAs targeting βII- and βIII-spectrin sequence (right). βII- and βIII-spectrin fluorescence intensities were assessed in spine necks, such as framed in bold white. Scale bars, 1 µm. **f,** Quantification of βII and βIII spectrins' fluorescence intensities in spine necks of control versus βII βIII double KO neurons. n = 237 and 210 spine necks from 3 independent experiments. Mann-Whitney test, ****: $p < 0.0001$.

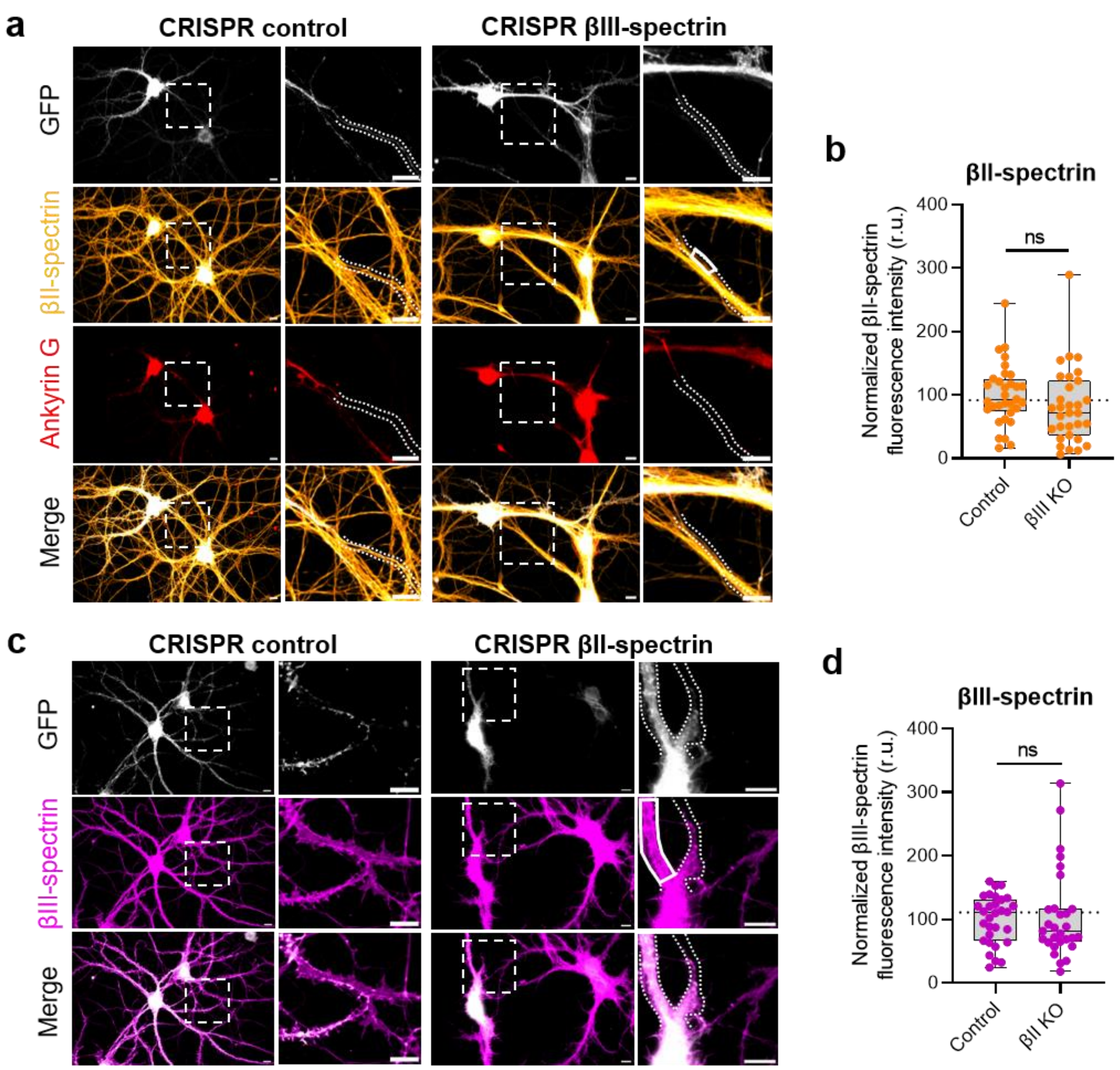


**Figure S4. CRISPR-Cas9 strategies targeting βII- or βIII-spectrin are specific. a,** Representative wide-field images of a non-transfected neuron (GFP-negative) alongside a neuron transfected with the CRISPR-Cas9 system and a gRNA control (left) or targeting a βIII-spectrin sequence (right). βII-spectrin fluorescence was assessed in axonal regions following the ankyrin G signal, such as framed in bold white. Scale bars, 10 µm. **b,** Quantification of βII-spectrin fluorescence intensity in distal axons of control versus βIII KO neurons. Each box represents the median with percentile. n = 32 and 31 neurons from 2 independent experiments. Mann-Whitney test, ns: non-significant. **c,** Representative wide-field images of a non-transfected neuron (GFP-negative) alongside a neuron transfected with the CRISPR-Cas9 system and a gRNA control (left) or targeting a βII-spectrin sequence (right). βIII-spectrin fluorescence was assessed in dendrites, such as framed in bold white. Scale bars, 10 µm. **d,** Quantification of βIII-spectrin fluorescence intensity in dendrites of control versus βII KO neurons. n = 30 and 29 neurons from 3 independent experiments. Mann-Whitney test, ns: non-significant

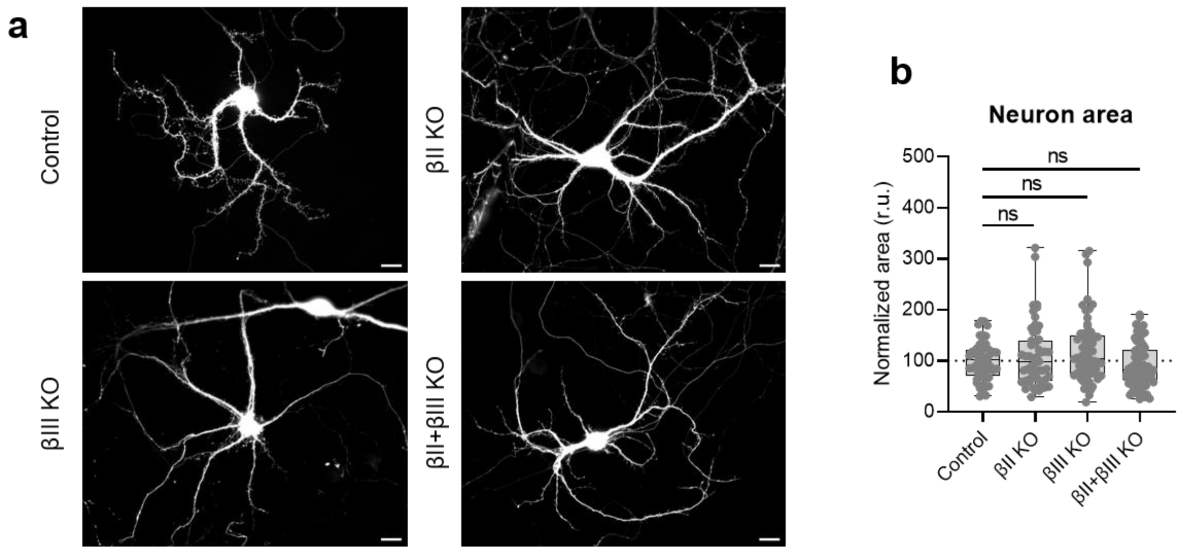


**Figure S5. Knocking-out βII- or βIII-spectrin at 8 DIV does not impair the total area of neurons. a,** Representative wide-field images of neurons transfected with the CRISPR-Cas9 and a gRNA control (left), targeting a βII-spectrin sequence (middle left), targeting a βIII-spectrin sequence (middle right), and after combining the two later constructs (right). Scale bars, 20 µm. **b,** Quantification of neuronal area based on the GFP fluorescence (reporter of transfection) in the different KO conditions. Each box represents the median with percentile. n = 50 to 72 neurons from 3 independent experiments. Mann-Whitney test, ns: non-significant.

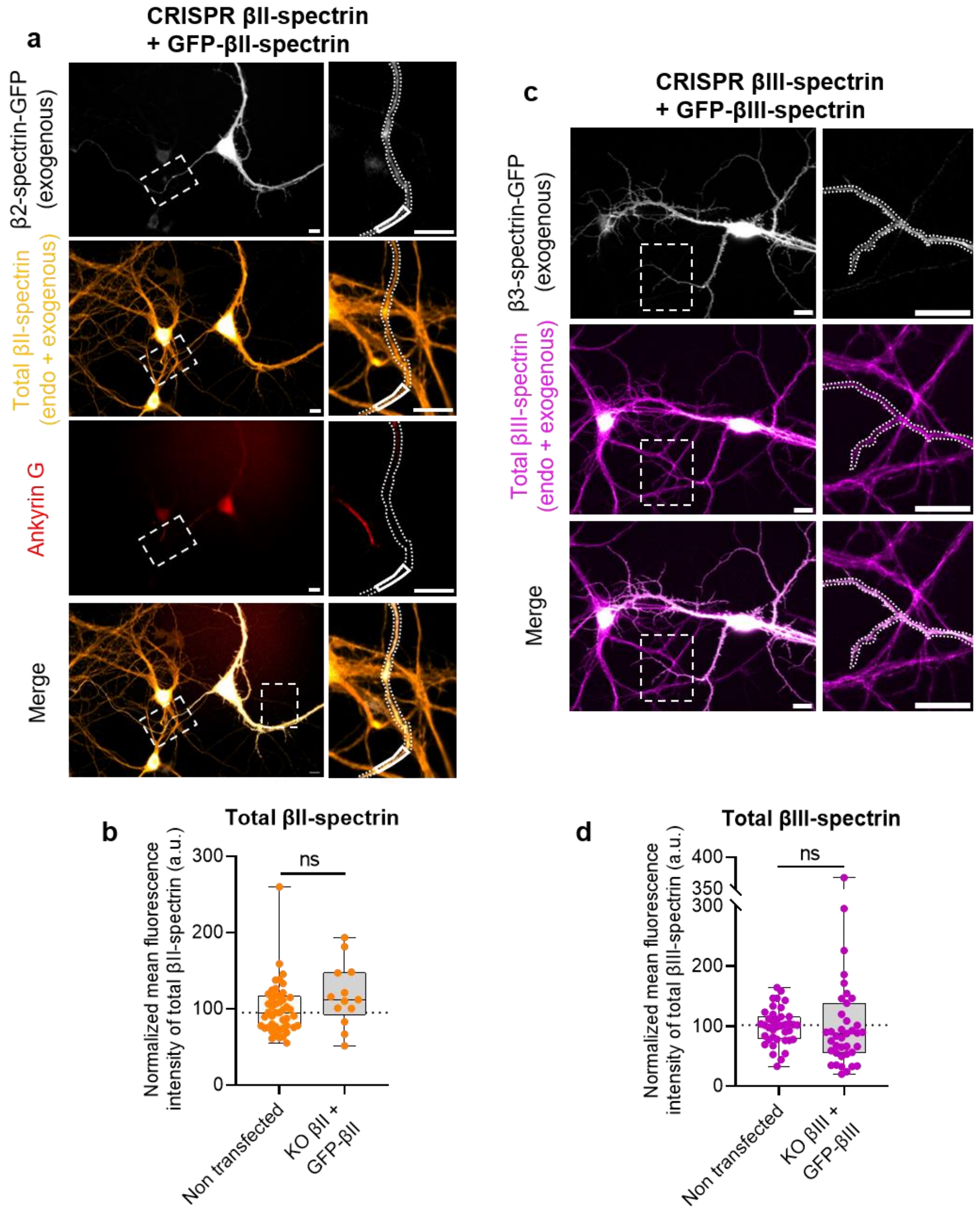


**Figure S6. The re-expression strategy allows near endogenous level tagged βII- or βIII-spectrin expression. a,** Representative wide-field images of a neuron co-transfected with the CRISPR-Cas9 system with a gRNA targeting a βII-spectrin sequence and a resistant plasmid for GFP-βII-spectrin expression. Total βII-spectrin was revealed by immunolabelling against βII-spectrin. Fluorescence was assessed in axonal regions following the ankyrin G signal, such as framed in bold white. **b,** Quantification of βII-spectrin fluorescence intensity in axons of non-transfected versus βII KO + rescue GFP-βII neurons. Each box represents the median with percentile. n = 13 to 48 neurons from 3 independent experiments. **c,** Representative wide-field images of a neuron co-transfected with the CRISPR-Cas9 system with a gRNA targeting a βIII-spectrin sequence and a resistant plasmid for GFP-βIII-spectrin expression. Total βIII-spectrin was revealed by immunolabelling against βIII-spectrin. Fluorescence was assessed in dendrites, such as framed in bold white. **d,** Quantification of βIII-spectrin fluorescence intensity in dendrites of non-transfected versus βIII KO + rescue GFP-βIII neurons. Each box represents the median with percentile. n = 39 neurons from 3 independent experiments. Mann-Whitney test, ns: non-significant, *: $p < 0.05$, **: $p < 0.01$, ***: $p < 0.001$.  Scale bars, 10 µm.

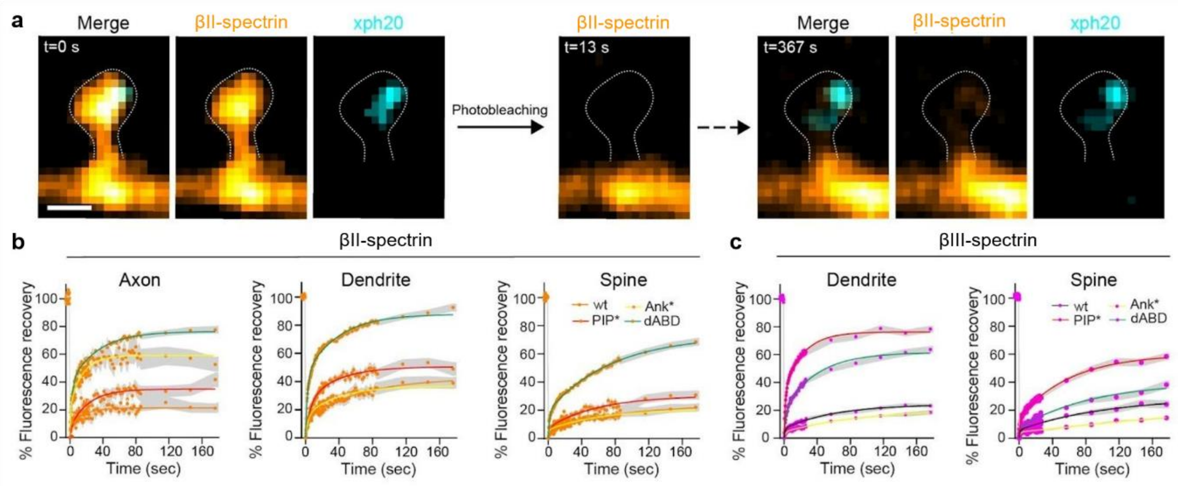


**Figure S7. FRAP measurements of βII- and βIII-spectrin. a,** Representative spinning disk images of a dendritic spine expressing GFP-βII-spectrin (orange) and an intrabody for PSD95 fused to mScarlett[76] (cyan) before photobleaching, just after photobleaching, and 5 minutes after photobleaching. Neurons were transfected with the CRISPR-Cas9 system with a gRNA targeting the βII-spectrin sequence and with a resistant GFP- βII-spectrin. Scale bar, 1 µm. **b,** Average FRAP curves of GFP-βII-spectrin and mutants (PIP*, Ank* and dABD) in axons, dendrites and spines. **c,** Average FRAP curves of GFP-βIII-spectrin and mutants (PIP*, Ank* and dABD) in dendrites and spines. All data are given as mean ± SEM.
